# Drivers of plankton community structure in intermittent and continuous coastal upwelling systems–from microscale in-situ imaging to large scale patterns

**DOI:** 10.1101/2023.05.04.539379

**Authors:** Moritz S Schmid, Su Sponaugle, Kelly R Sutherland, Robert K Cowen

## Abstract

Eastern Boundary Systems support major fisheries whose early life stages depend on upwelling production. Upwelling can be highly variable at the regional scale, with substantial repercussions for new productivity and microbial loop activity. A holistic assessment of plankton community structure is challenging due to the range in body forms and sizes of the taxa. Thus, studies that integrate the classic trophic web based on new production with the microbial loop are rare. Underwater imaging can overcome this limitation, and together with machine learning, enables fine resolution studies spanning large spatial scales. We used the In-situ Ichthyoplankton Imaging System (ISIIS) to investigate the drivers of plankton community structure in the northern California Current, sampled along the Newport Hydrographic (NH) and Trinidad Head (TR) lines, in OR and CA, respectively. The non-invasive imaging of particles and plankton (250μm –15cm) over 1644km (30 transects) in the winters and summers of 2018 and 2019 yielded 1.194 billion classified plankton images. The imaged plankton community ranged from protists, crustaceans, and gelatinous taxa to larval fishes. To assess community structure, >2000 single-taxon distribution profiles were analyzed using high resolution spatial correlations. Co-occurrences on the NH line were consistently significantly higher off-shelf while those at TR tended to be highest on-shelf. Taxa co-occurrences at TR increased significantly with upwelling strength and in 2019 TR summer co-occurrences were similar to those on the NH line. Random Forests models identified the concentrations of microbial loop taxa such as protists, *Oithona* copepods, and appendicularians as important drivers of co-occurrences at NH line, while at TR, cumulative upwelling and chlorophyll a were of the highest importance. Our results indicate that the microbial loop is actively driving plankton community structure in intermittent upwelling systems such as the NH line and may induce temporal stability. Where upwelling is more continuous such as at TR, primary production may dominate patterns of community structure, obscuring the underlying role of the microbial loop. Future changes in upwelling strength are likely to disproportionately affect plankton community structure in continuous upwelling regions, while high microbial loop activity enhances community structure resilience.

## 1 Introduction

Community structure in an ecological system is defined as the interactions among organisms within the community (Verity and Smetacek, 1996; Smetacek, 2012; Lima-Mendez et al., 2015). Planktonic community structure in the ocean and the processes driving it determine energy transfer through the trophic web, setting up fisheries and top predators (Brown et al., 2004), as well as carbon sinks. Community structure can be assessed through the lens of taxa co-occurrence. As co-occurrence is driven by processes enabling coexistence within an ecosystem, such as niche separation (MacArthur, 1958; Chesson et al., 2000; Lindegren et al., 2020), co-occurrences together with their biotic and abiotic environmental envelope describe ecologically important patterns that enable the investigation of community structure (HilleRisLambers et al., 2012; Williams et al., 2014; Ríos-Castro et al., 2022).

The oceanic environmental envelope is changing (Bakun et al., 1990; Doney et al., 2012; Bakun et al., 2015; Bograd et al., 2022). While climate change is a global phenomenon, key systems in which climate change has particularly strong effects are Eastern Boundary Upwelling Systems (EBUSs), due to the disproportional contribution of EBUSs to global ocean productivity and ecosystem services such as fisheries (Bograd et al., 2022). One such EBUS is the California Current Ecosystem which extends from British Columbia, Canada, to Baja California Sur in Mexico and exhibits strong physical and ecosystem variability on seasonal, interannual, and decadal time scales (Ware and Thomson, 2005; Barth et al., 2007; Checkley and Barth, 2009). The Northern California Current (NCC), extends from the northern border of the California Current Ecosystem southward to Cape Mendocino, CA, and encompasses variable oceanography.

Upwelling in the NCC varies with latitude and season, with distinct downwelling (winter) and upwelling (spring-summer) seasons off OR and WA contrasted with persistent, stronger upwelling off northern CA (Bograd et al., 2009; García-Reyes and Largier, 2012). During upwelling, cold, nutrient- and CO_2_-rich waters reach the euphotic zone (Barth et al., 2005; Kirincich et al., 2005; Hales et al., 2006), fueling high phytoplankton production (Dickson and Wheeler, 1995; Hales et al., 2006). Spring-summer upwelling typically occurs in intermittent events (3-10 d) and is demarcated by brief relaxation periods that cumulatively fuel strong primary and secondary production in the system (Hickey and Banas, 2003; Feinberg and Peterson, 2003; Shaw et al., 2010). The NCC shelf in mid and northern Oregon (e.g., Newport at 45°N) is relatively wide, allowing for higher retention of upwelled waters compared to southern OR and northern CA locations such as Cape Blanco (42.8°N) with a narrower shelf. Circulation tends to closely track bathymetry (Lentz and Chapman, 1989; Kirincich et al., 2005; Hickey and Banas, 2008), with the coastal upwelling jet meandering off the shelf south of Heceta Bank (Barth et al., 2000).

Tightly woven into the marine food web is the microbial loop, which enhances water column recycling of carbon and nutrients, making these available again to higher trophic levels (Turner et al., 2015; Cavan et al., 2019). While the microbial loop is often associated with low-latitude marine ecosystems with low nutrient levels and more recycled production (Azam et al., 1983), the microbial loop is ever-present, even in temperate areas where upwelling is prevalent (Wilkerson et al., 1987; González et al., 2004). In intermittent upwelling regimes, smaller plankters associated with the microbial food web can become dominant (Mousseau et al., 1998). Though patterns and processes of microbial cycling have been extensively studied (Pomeroy, 1974; Azam et al., 1983; Kirchman, 2010), its influence is rarely examined beyond lower trophic levels, and it is often ignored in upwelling systems where the primary focus has been on new production and classical food chains (but see Vargas et al., 2007).

The central importance of upwelling to the NCC comes with some negative consequences in the form of hypoxic and anoxic events. When low oxygen water is upwelled onto the shelf and phytoplankton blooms collapse, bottom water can quickly become depleted of oxygen (Chan et al., 2008; 2019). Increasingly frequent, such events are also associated with low pH (ocean acidification) conditions (Feely et al., 2008; Chan et al., 2019), both having significant negative effects on demersal habitats and organisms (Doney et al., 2020; Nagelkerken and Connell, 2022). Upwelling regimes are poised to shift as a result of changes in wind forcing due to global climate change (Bakun, 1990; Bakun et al., 2015; Buil et al., 2021). The resulting poleward intensification of upwelling and equatorward reduction in upwelling will most certainly affect new productivity and thus activity of the microbial loop. In particular, the northern NCC is predicted to experience more upwelling-favorable winds in the future (Buil et al., 2021).

Simultaneously, climate change has the potential to affect oceanographic processes at all spatial scales that comprise the environmental envelopes experienced by marine taxa. Effects are likely to be evident in taxa distributions, community composition, and community structure at scales ranging from microscale (e.g., predator-prey interactions, nutrient uptake in phytoplankton), fine scale (e.g., plankton thin layers and internal waves), sub-mesoscale (e.g., coastal processes such as cross-shore transport and upwelling), mesoscale (e.g., eddies, wind stress curl), and large basin scale [e.g., marine heat waves, Pacific Decadal Oscillation (PDO); Bakun and Nelson, 1991; Denman and Gargett, 1995; Mantua and Hare, 2002; Dickey and Bidigare, 2005; Prairie et al., 2012].

Historically, net-based plankton sampling has not adequately resolved planktonic communities at the micro-, and fine scales that are important for plankton dynamics (Haury et al., 1978; Yamazaki et al., 2002; Benoit-Bird et al., 2013; Schmid et al., 2019; Robinson et al., 2021). In response, in-situ imaging instruments have been developed over the past few decades (Ortner et al., 1979) that can overcome this limitation. Today a variety of systems exist that have been designed for specific tasks: for instance, UVP6 (Picheral et al., 2022), Zooglider (Ohman et al., 2019), PlanktonScope (Song et al., 2020), the Scripps Plankton Camera System (Orenstein et al., 2020), and the In-situ Ichthyoplankton Imaging System (ISIIS; Cowen and Guigand, 2008). Advantages of imaging systems include their non-destructive and high spatial resolution sampling capability (Lombard et al., 2019) as well as efficient imaging of plankton traits (Schmid et al., 2018; Vilgrain et al., 2021; Lertvilai and Jaffe, 2022). Data from imaging systems are often analyzed using machine learning due to the volume of data generated (Luo et al., 2018; Irisson et al., 2021). Together with additional onboard sensors (e.g., fluorometers, oxygen probes, CTDs) these imaging systems can describe the plankton community and their environmental envelope with high spatial resolution, providing new insight into plankton community structure (Briseño-Avena et al., 2020; Robinson et al. 2021).

To investigate the drivers of planktonic community structure in the NCC ecosystem, we deployed the ISIIS along two cross-shelf transects that varied in their seasonal patterns of upwelling. Sampling across two seasons (winter and summer) for two years (2018, 2019), in conjunction with a deep learning data pipeline, yielded a very large dataset for examining plankton community structure. To obtain a holistic view of community structure we used a spatially explicit high-resolution correlation of taxa distributions. The co-occurrence of a wide range of organisms spanning from primary producers and protists, through gelatinous plankton and crustacean zooplankton, to larval fishes, in the context of their biotic and abiotic environment was used to disentangle the degree to which community structure is driven by upwelling strength and new productivity versus the potential impact of the microbial loop. With climate change beginning to affect the California Current Ecosystem, it is important to identify current drivers of plankton community structure such that we can better anticipate future changes to the structure of the water column that may disrupt the costal marine food web including valuable fisheries.

## 2 Materials and Methods

### 2.1 Study area

Thirty transects ranging between 24 and 86 km in length were sampled along the Newport Hydrographic (NH) line as well as the Trinidad Head (TR) line during the winters (February-March) and summers (July-August) of 2018 and 2019. Located off Newport, Oregon (Fig. 1), the NH Line has been sampled since 1961 (Peterson and Miller, 1975), while the TR line off northern California has been sampled since 2007 (Robertson and Bjorkstedt, 2020). Both transects are part of regular net-based sampling efforts by the National Oceanic and Atmospheric Administration (NOAA) with a focus on determining the plankton community structure and the biophysical drivers of the recruitment of commercially important fishes. Imagery data were collected during both day and night hours, with daytime transects commencing at least 1h after sunrise and ending at least 1h before sunset, and nighttime transects commencing at least 1h after sunset and ending at least 1h before sunrise.

**Figure 1.**
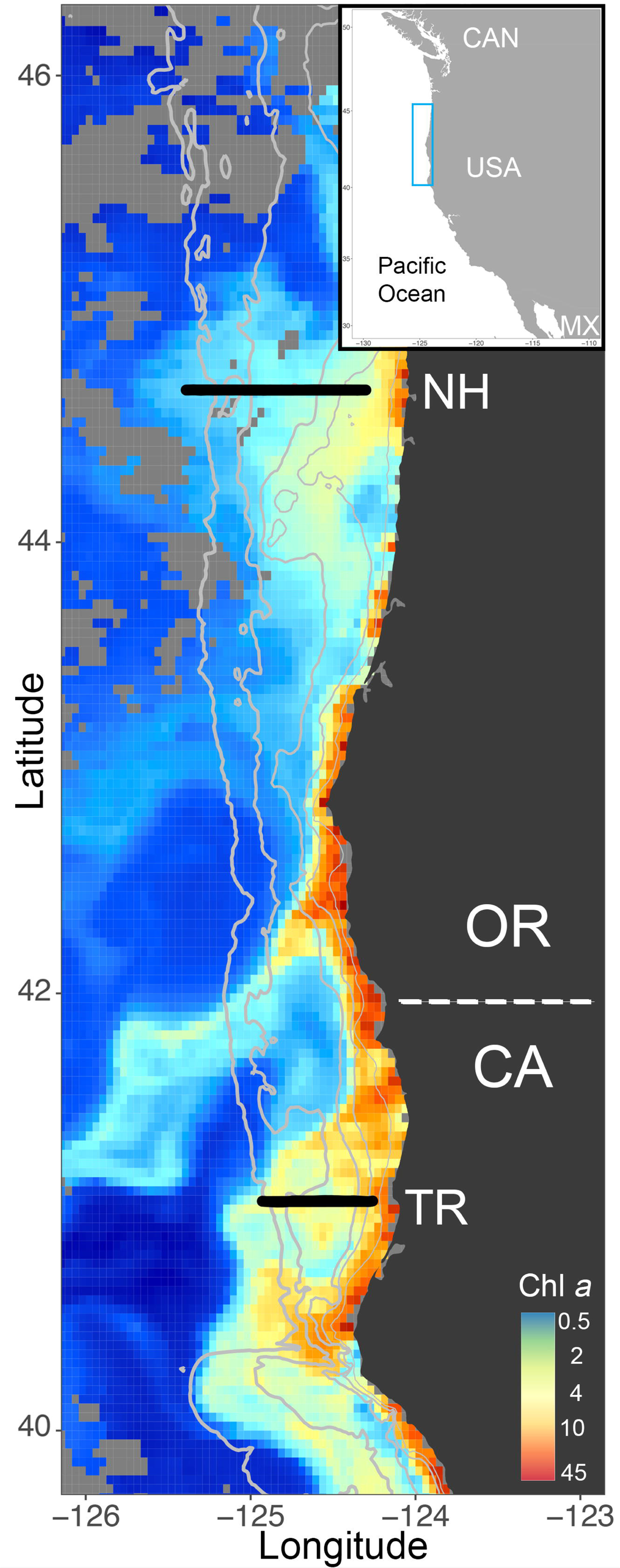
ISIIS transects along the Newport Hydrographic (NH) and Trinidad Head (TR) lines (black solid line) where sampling occurred in winter and summer 2018 and 2019. Chlorophyll *a* from Aqua Modis ocean color on July 10, 2018 shows the often higher productivity in the northern California part of the Northern California Current, where the shelf is narrower than farther north at the NH Line (depth contours in solid grey lines at 50m, 100m, 200m, 1000m, 2000m). Light grey pixels indicate non-available data from Aqua Modis.

### 2.2 In-situ Ichthyoplankton Imaging System (ISIIS)

ISIIS (Cowen and Guigand, 2008) is a towed shadowgraph and line-scan imaging system, that scans a large volume of water (150 -185 L^−1^) to quantitatively sample abundant meso-zooplankton as well as rarer ichthyoplankton (Cowen et al., 2013). ISIIS’s large imaging frame, with a 13 × 13-cm field of view and 50 cm depth of field allows for the undisturbed imaging of a variety of plankton taxa including fragile gelatinous zooplankton (McClatchie et al., 2012; Luo et al., 2014). The resulting images have a pixel resolution of 66 μm and are recorded as continuous videography. Data are sent to a top-side computer using a fiber optic cable where ISIIS data are time-stamped. ISIIS is equipped with a CTD (Sea-Bird SBE 49 FastCAT), as well as a dissolved oxygen probe (Sea-Bird 43), fluorescence sensor (Wet Labs FLRT), and photosynthetically active radiation sensor (PAR; Biospherical QCP-2300). ISIIS is towed behind the ship at 2.5 m s^-1^ where it undulates on each cross-shelf transect between 1 m and 100 m depth or as close as 2 m above the seafloor in shallower water. ISIIS has been used in various ecosystems with differing scientific objectives, such as the investigation of larval fish distributions at eddy fronts (Schmid et al., 2020) and fine-scale plankton patchiness in the Straits of Florida (Robinson et al., 2021), larval fish distributions in the context environmental gradients in the NCC (Swieca et al., 2020; Briseño-Avena et al., 2020), the investigation of zooplankton individual-level interactions and parasitism in the Gulf of Mexico (Greer et al., 2021), and cross-ecosystem comparisons of a gelatinous grazer (Greer et al., 2023).

### 2.3 Sparse convolutional neural net

ISIIS imagery data were processed following Luo et al. (2018) and Schmid et al. (2020), with a full open-sourced, pipeline code (Schmid et al., 2021). After the collected video data were flat-fielded and segmented into single regions of interest (ROIs; i.e., a single plankton specimen) using a k-harmonic means clustering algorithm, a training library of images was created by choosing representative images from all 2018 and 2019 transects. The training library contained 82,909 images spanning 170 different classes, ranging from protists and phytoplankton to larval fishes. The sCNN (SparseConvNets with Fractional MaxLJPooling; Graham et al., 2015; Luo et al. 2018) was trained until the error rate plateaued at ∼ 5% after 399 epochs.

The 170 original classes in the training library were mapped onto 67 broader groups (e.g., chaetognaths of different shapes merged into one group). After removing five different unknown groups, 62 taxonomic groups remained for ecological analyses. A random subset of images was classified by two human annotators and used for probability filtering following Faillettaz et al. (2016), an approach that removes very low probability images from the dataset, achieving 90% predictive accuracy per taxon. Removal of these “low confidence images” still allows for the prediction of true spatial distributions (Faillettaz et al., 2016). An independent subsample of the remaining images was again classified by the same two human annotators and the results compared with the automated classification. The resulting confusion matrix was used to calculate taxon-specific correction factors:

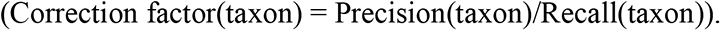

Individuals and environmental data were binned into 1 m vertical strata and plankton concentrations (ind. m^-3^) estimated based on the volume of imaged seawater. Plankton concentrations were then adjusted by applying the taxon-specific correction factors.

### 2.4 Environmental and ecological data analyses

To estimate the upwelling strength on each transect, we calculated the cumulative daily Coastal Upwelling Transport Index (CUTI^1^; Jacox et al., 2018) for the 10 d prior to sampling of a transect. This period was selected to account for the lag between physical forcing (i.e., nutrient upwelling) and phyto-(∼7 d) and zooplankton (∼13-16 d) abundances (Spitz et al., 2005). To encompass plankton ranging from phyto-to zooplankton, we selected an intermediate lag of 10 d (Swieca et al., *in review*). Plankton community structure was assessed in two approaches using spatially explicit Spearman rank correlations as measure of the co-occurrence of taxa.

### 2.5 Low spatial resolution taxa co-occurrence

To better describe the environmental drivers of species distributions, transects were divided into an on-shelf portion and off-shelf portion based on the longitude of the 200 m isobath. Spearman rank correlations between the concentrations of each taxon and the remainder of the plankton community were calculated for both portions of a transect, with the underlying data binned vertically in 1-m strata (Fig. 2). Thus, each taxon was correlated 122 times per transect: 61 correlations with the other taxa on the shelf, and 61 correlations with other taxa off the shelf. Correlations were computed for all possible combinations of the 62 taxa along the 30 transects and only correlations that were significant at p < 0.01 were used for further analyses.

**Figure 2.**
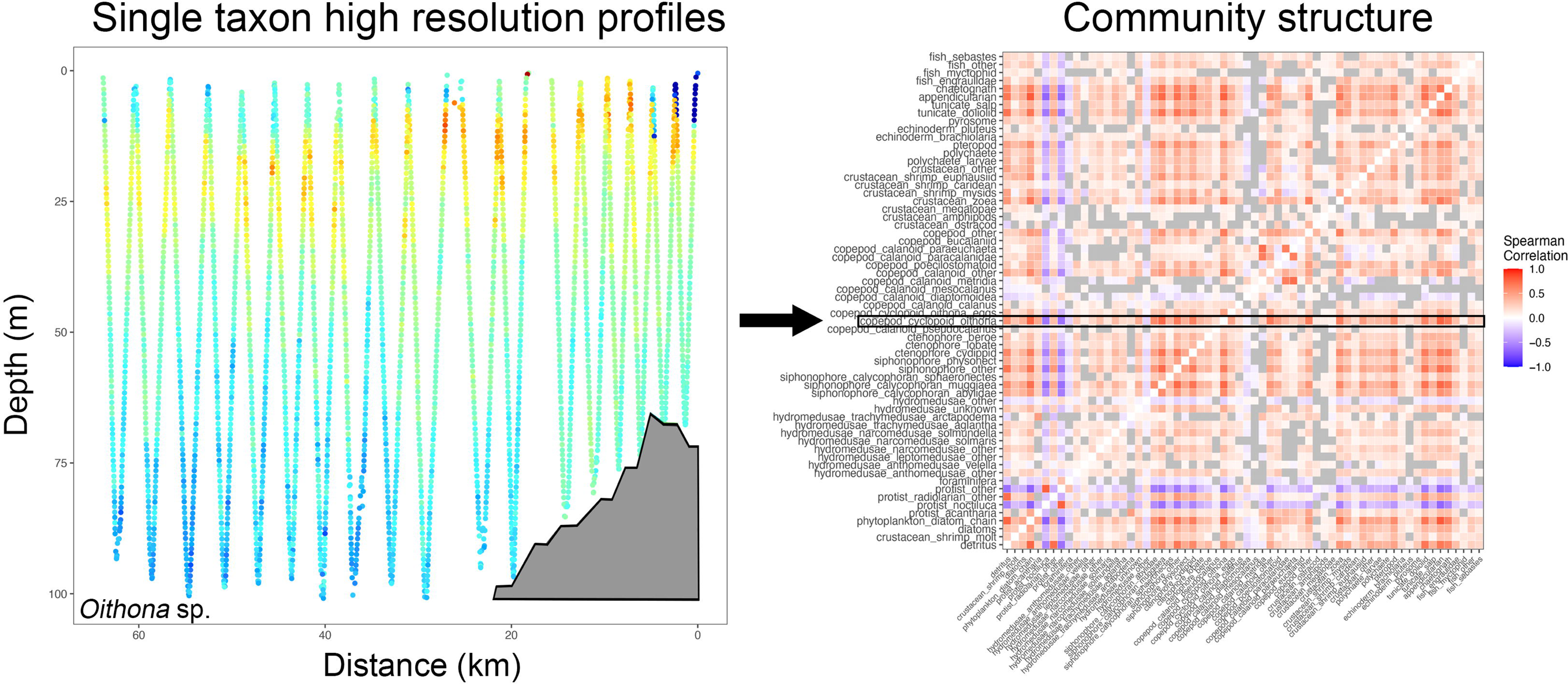
High spatial distribution profiles (e.g., here *Oithona* sp., left panel – shelf delineated as a grey polygon) are correlated with all other taxa and led to correlograms that depict co-occurrence amongst taxa (right panel). In this example, the on-shelf portion of the left panel would result in one correlogram, and the off-shelf part in another. The black box shows *Oithona* sp. co-occurrence with other taxa (blue hues indicating negative correlation and red hues positive correlation, grey boxes are correlations that are not significant at p <0.01), while other taxa correlations above and below depict correlations of the remainder of the plankton community on the transect.

Co-occurrences were compared between the two years, seasons, study sites, and shelf condition (on- and off-shelf). To determine whether on-shelf and off-shelf co-occurrences differed among the different years, seasons, sites, and shelf conditions, we used Wilcoxon Rank Sum tests with a confidence level of 0.999. To delineate patterns of taxa co-occurrences, coefficients of variation of the co-occurrences were calculated for each possible year, season, site, and shelf condition combination. Finally, to test for differences between NH and TR, the co-occurrences were regressed against CUTI at the two different study sites using a Wilcoxon Rank Sum test.

### 2.6 High spatial resolution taxa co-occurrence and their environmental drivers

Based on the original 1-m vertically stratified data, mixed layer depth (Kara et al., 2000), Brunt Vaisala Frequency, and geostrophic dynamic height anomalies (both using ‘gsw’ R package which follows TEOS-10 definitions) were calculated along each transect to account for mixing depth, stratification strength, and influence of the upwelling front, respectively. The data were then projected onto a 10-m vertically stratified grid while keeping the underlying 1 m vertical data structure intact. Spearman rank correlations were calculated similarly to 2.4.1, but instead of calculating one correlation for a taxon per on- or off-shelf, the 10 m grid was used to calculate a correlation for each 10 m interval of the water column. This enabled the retention of both vertically stratified environmental and co-occurrence data. Only Spearman rank correlations that were significant at p < 0.01 were used in further analyses.

Resulting environmental and taxon-specific data (Table 1) were used in two Random Forests models, one for each of the two transect lines. In each case the modeled response variable was the correlation coefficient for a specific 10-m vertical section of the grid; however, all data collected on NH or TR were combined for the most generalist model. Random Forest analysis (Breiman et al., 2001) was carried out using the ‘caret’ package in R (Kuhn et al., 2008), in the ‘ranger’ RF implementation, and variable importance was assessed based on permutation importance. Partial dependence plots were used to investigate the specific non-linear effects of the 10 most important explanatory variables per model.

**Table 1:** Variables used in Random Forests modeling included biotic and abiotic environmental variables (1-14) and taxa concentrations derived from underwater imaging (15-83).

| Name | Unit | Name | Unit |
| --- | --- | --- | --- |
| 1 Shelf (On/Off) | - | 43 detritus | ind. m<sup>-3</sup> |
| 2 Distance | km | 44 diatoms | " |
| 3 Depth | m | 45 echinoderm_brachiolaria | " |
| 4 Year (2018 / 2019) | - | 46 echinoderm_pluteus | " |
| 5 Season (winter / summer) | - | 47 fish_clupeiformes | " |
| 6 Cumulative CUTI (10 day) | m<sup>-3</sup> s<sup>-1</sup> | 48 fish_flatfish | " |
| 7 Temperature | °C | 49 fish_long_slender | " |
| 8 Salinity | - | 50 fish_myctophid | " |
| 9 Oxygen | ml l<sup>-1</sup> | 51 fish_sebastes | " |
| 10 Chlorophyll *a* | µg l<sup>-1</sup> | 52 fish_unknown | " |
| 11 Density | kg m<sup>-3</sup> | 53 foraminifera | " |
| 12 Geostrophic Dynamic Height Anomaly | m<sup>2</sup> s<sup>-2</sup> | 54 hydromedusae_anthomedusae_other | " |
| 13 Mixed layer depth | m | 55 hydromedusae_anthomedusae_velella | " |
| 14 Brunt Vaisala Frequency squared | rad<sup>2</sup> s<sup>-2</sup> | 56 hydromedusae_leptomedusae_eutonia | " |
| 15 appendicularians | ind. m<sup>-3</sup> | 57 hydromedusae_leptomedusae_clytia_mitrocoma | " |
| 16 chaetognaths | " | 58 hydromedusae_leptomedusae_other | " |
| 17 copepod_calanoid_calanus | " | 59 hydromedusae_narcomedusae_aegina | " |
| 18 copepod_calanoid_diapomoida | " | 60 hydromedusae_narcomedusae_other | " |
| 19 copepod_calanoid_mesocalanus | " | 61 hydromedusae_narcomedusae_solmaris | " |
| 20 copepod_calanoid_metridia | " | 62 hydromedusae_narcomedusae_solmundella | " |
| 21 copepod_calanoid_other | " | 63 hydromedusae_other | " |
| 22 copepod_calanoid_paracalanidae | " | 64 hydromedusae_trachymedusae_aglantha | " |
| 23 copepod_calanoid_paraeuchaeta | " | 65 hydromedusae_trachymedusae_aglaura | " |
| 24 copepod_calanoid_pseudocalanus | " | 66 hydromedusae_trachymedusae_arctapodema | " |
| 25 copepod_carcass | " | 67 hydromedusae_unknown | " |
| 26 copepod_cyclopoid_oithona | " | 68 phytoplankton_diatom_chain | " |
| 27 copepod_cyclopoid_oithona_eggs | " | 69 polychaete | " |
| 28 copepod_eucalaniid | " | 70 polychaete_larvae | " |
| 29 copepod_other | " | 71 protist_acantharia | " |
| 30 copepod_poecilostomatoid | " | 72 protist_noctiluca | " |
| 31 crustacean_amphipods | " | 73 protist_other | " |
| 32 crustacean_megalopae | " | 74 protist_radiolarian_other | " |
| 33 crustacean_ostracod | " | 75 pteropod | " |
| 34 crustacean_other | " | 76 pyrosome | " |
| 35 crustacean_shrimp_caridean | " | 77 siphonophore_calycophoran_abyldae | " |
| 36 crustacean_shrimp_euphausiid | " | 78 siphonophore_calycophoran_muggiaea | " |
| 37 crustacean_shrimp_molt | " | 79 siphonophore_calycophoran_sphaeronectes | " |
| 38 crustacean_shrimp_mysids | " | 80 siphonophore_other | " |
| 39 crustacean_zoea | " | 81 siphonophore_physonect | " |
| 40 ctenophore_beroe | " | 82 tunicate_doliolid | " |
| 41 ctenophore_cydippid | " | 83 tunicate_salp | " |
| 42 ctenophore_lobate | " |  |  |

## 3 Results

1.194 billion plankton images were classified based on 195 h of underwater imagery, traversing a total of 1644 km along 30 transects ranging from 24-86 km (Supplementary Material Table S1). The vast diversity of NCC plankton imaged included taxa such as appendicularians, crab zoea and megalopa, different types of copepods as well as hydromedusae, pteropods, chaetognaths, ctenophores, salps, several groups of larval fish (Fig. 3, Supplementary Material Table S1), among others. Dense thin layers of different plankton taxa were observed frequently during deployments and analysis of the imagery showed these dense layers consisted of > 25,000 calanoid copepods per m^-3^ (Feb 2018 on the TR line, Supplementary Material Table S1), or of >1,300 crab zoea per m^-3^ (Feb 2018 on the NH line). Thin dense layers of doliolids reached densities of >11,000 individuals per m^-3^ (e.g., NH line in July 2018), while lobate ctenophores accumulated to >800 individuals per m^-3^ (March 2019 on the NH line). Appendicularian accumulations of >10,000 individuals per m^-3^ were found on the TR line in July 2019.

**Figure 3.**
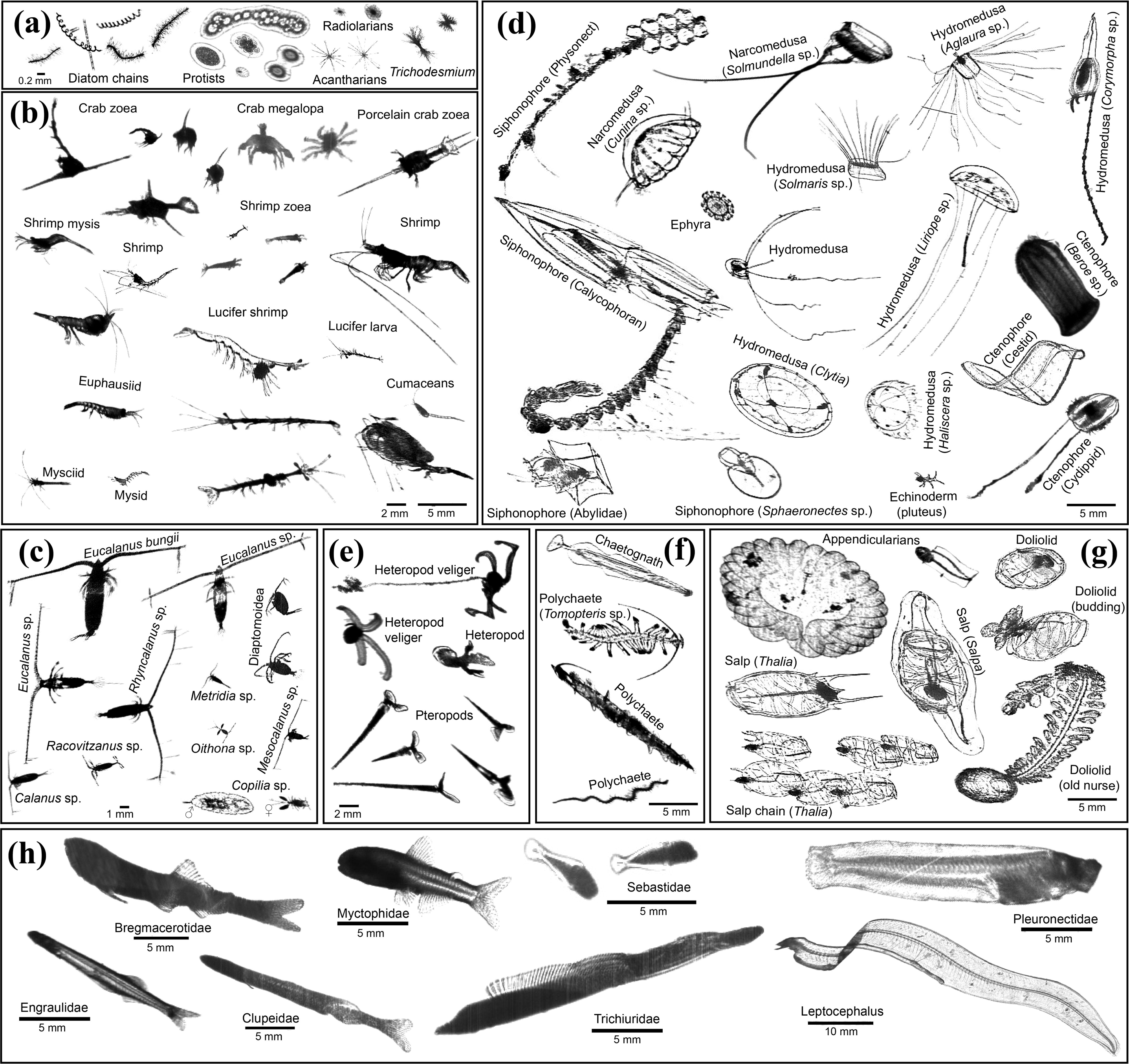
ISIIS images of key taxa in the northern California Current. (A) Primary producers and protists; (B-C) crustaceans (D) cnidarians, ctenophores, and echinoderms; (E) heteropods and pteropods; (F) chaetognaths and polychaetes; (G) pelagic tunicates; (H) larval fishes.

Comparison of the environmental conditions at the NH and TR lines revealed distinct upwelling signatures in seawater densities along both sampling lines in the summers of 2018 and 2019 (see representative transects in Figs. 4, 5). Winter upwelling was less evident in TR line pycnoclines, but more so in cumulative CUTI upwelling 10 d prior to sampling (Fig. 6). Sea surface temperatures on the NH line peaked at 17.5°C in 2018, while water on the TR line remained substantially cooler. Surface salinities at the NH line were relatively fresh at 30, while most of the water column on the TR line was > 32.5. Oxygen levels fell to < 2 ml l^-1^ levels on the NH line in summer 2018, coinciding with substantial upwelling. Chlorophyll *a* was highest on the TR line in summer 2018 with levels reaching up to 45 μg l^-1^. The temperature profile along the NH line in summer 2019 closely mimicked that of 2018, with surface temperatures > 17.5 °C (Fig. 5). In 2019, oxygen levels fell to < 3 ml l^-1^ in summer in near-bottom areas on the shelf, while sub-surface chlorophyll maxima reached 7.5 μg l^-1^ at both sampling sites in summer.

**Figure 4.**
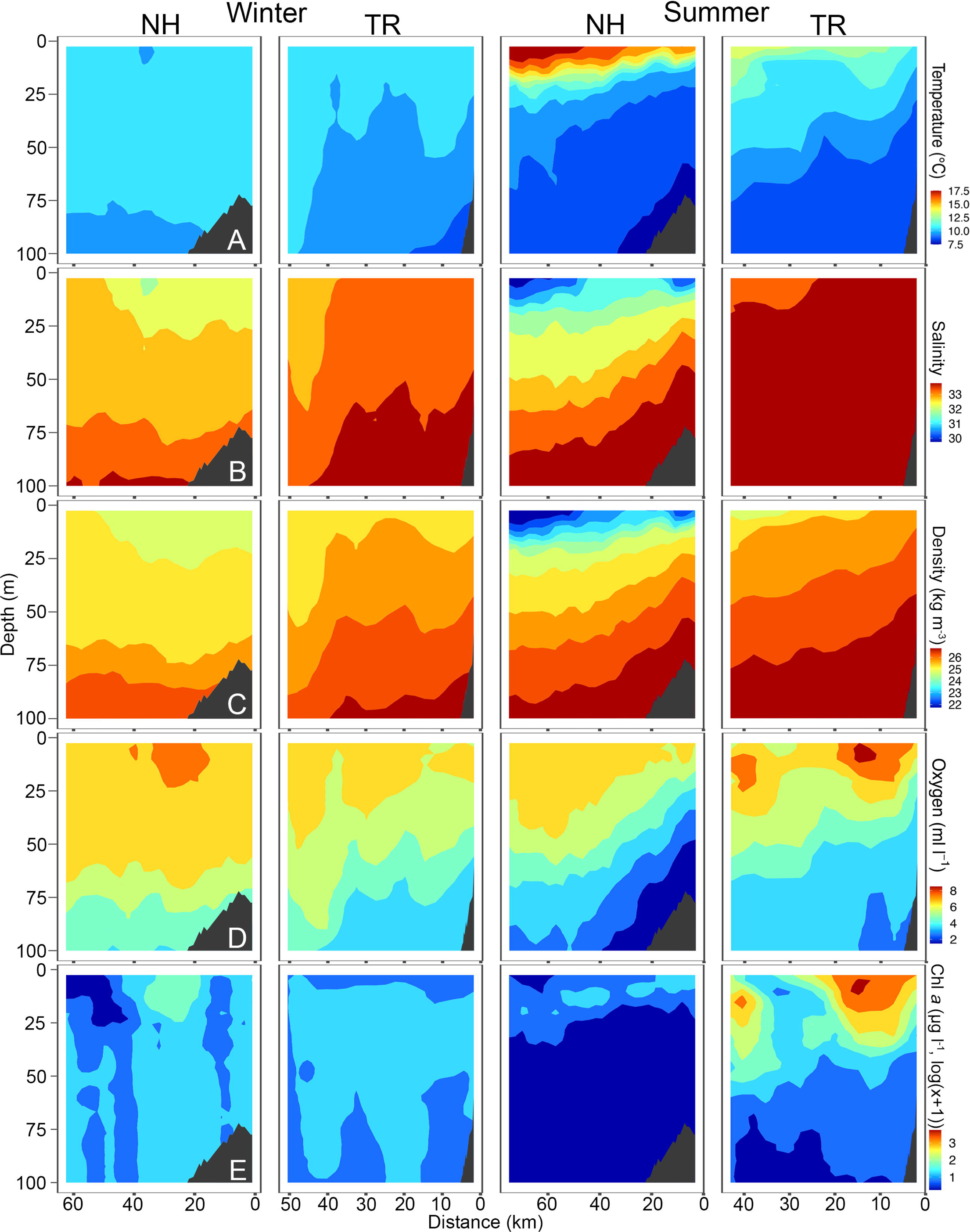
Temperature (A), Salinity (B), Density (C), Oxygen (D) and Chlorophyll *a* (chl *a*, E) across the Newport Hydrographic (NH) and Trinidad Head (TR) transects in winter and summer 2018. Winter sampling on the NH and TR shown here was carried out February 16 and 21, respectively, while summer sampling was carried out on July 10 and 7, respectively. Note that Chl *a* is plotted in log(x+1) due to the values ranging from 0.01 to 45 μg l^-1^. Shelf indicated in dark grey.

**Figure 5.**
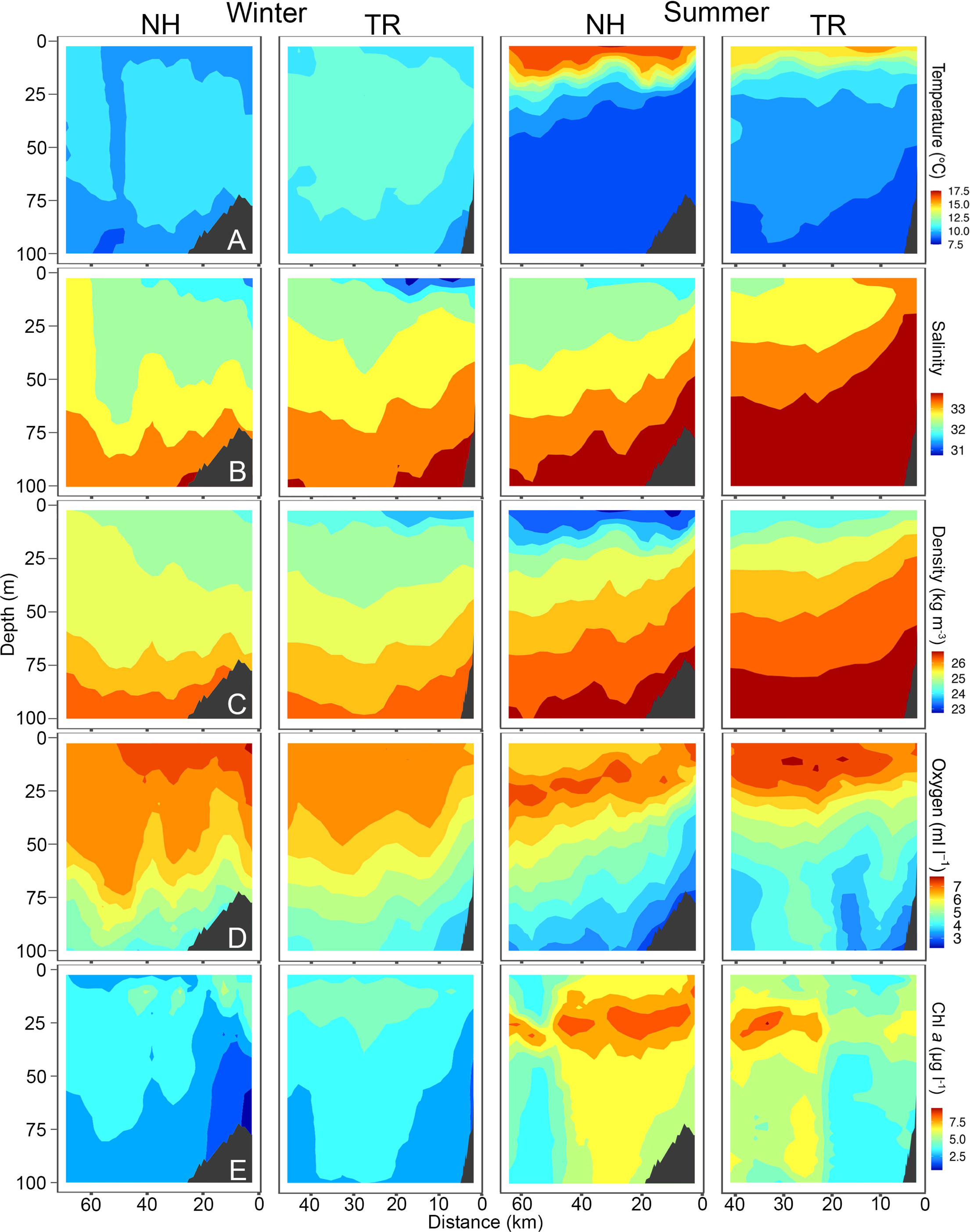
Temperature (A), Salinity (B), Density (C), Oxygen (D) and Chlorophyll *a* (chl *a*, E) across the Newport Hydrographic (NH) and Trinidad Head (TR) transects in winter and summer 2019. Winter sampling on the NH and TR shown here was carried out on March 6 and 8, respectively, while summer sampling was carried out on July 23 and 18, respectively. Shelf indicated in dark grey.

**Figure 6.**
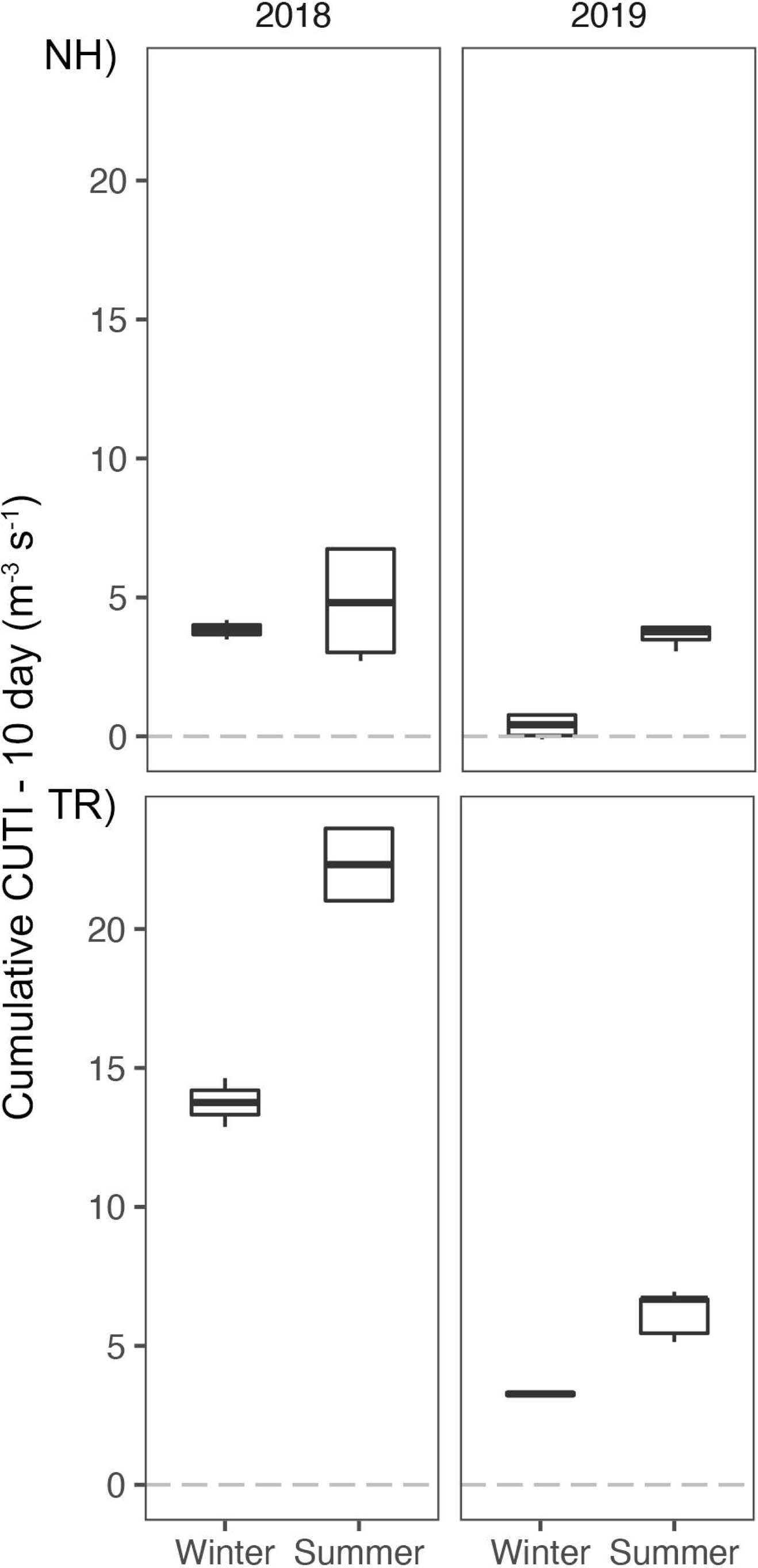
Cumulative Coastal Upwelling Transport Index (CUTI) over 10 d prior to sampling at the Newport Hydrographic (NH) and Trinidad Head (TR) lines.

Cumulative CUTI upwelling 10 d prior to sampling each transect was always higher on the TR line relative to the NH line (Fig. 6). The mean 10-day CUTI for the NH line during winter 2018 was 3.8 m^-3^ s^-1^, while it was 4.8 m^-3^ s^-1^ during summer (Fig. 6). Upwelling in 2019 was markedly lower, with a mean of 0.4 m^-3^ s^-1^ during winter and 3.6 m^-3^ s^-1^ during summer. On the TR line, 2018 10-d CUTI upwelling reached 13.8 m^-3^ s^-1^ and 22.3 m^-3^ s^-1^ during winter and summer, respectively, while upwelling in 2019 followed the NH trend and was reduced to 3.3 m^-3^ s^-1^ and 6.2 m^-3^ s^-1^ during winter and summer, respectively.

TS-diagrams demonstrate that temperatures on the NH line in summer were consistently the highest ones measured during the study (Fig. 7). Winter profiles were characterized by a substantially smaller water temperature range on both transects. While warm summer surface waters on the NH line were also the freshest found at either site, winter water was fresher on the TR line than on the NH line. Winter and summer water at both sampling sites showed a distinct seasonal signal (Fig. 7).

**Figure 7.**
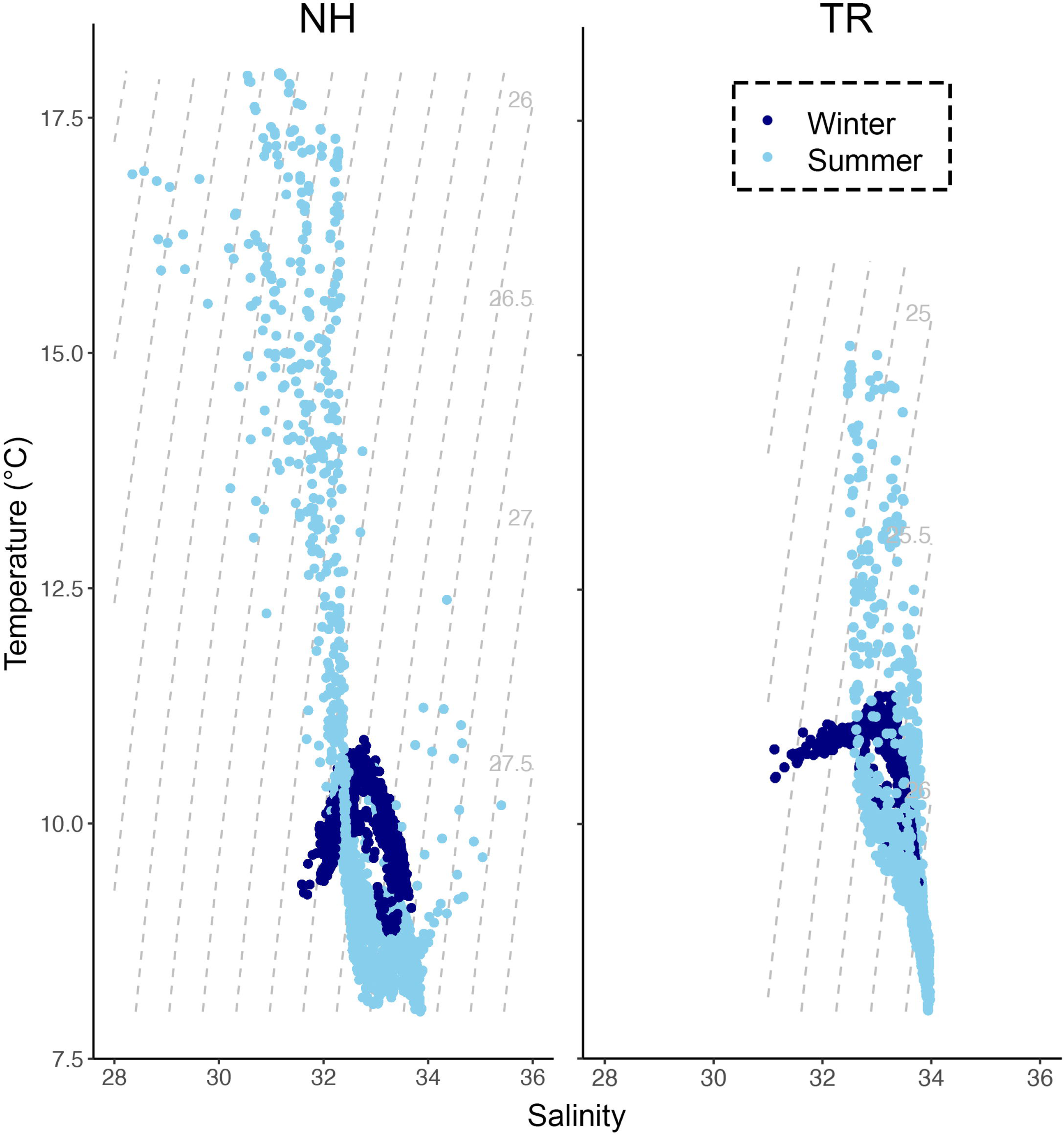
T-S diagrams during summer and winter sampling (2018, 2019) along the Newport Hydrographic (NH), and Trinidad Head (TR) lines. Light grey numbers and dotted lines indicate isopycnals.

### 3.1 Low spatial resolution taxa co-occurrences

Co-occurrences among taxa ranged from strongly negatively correlated at -1 to strongly positively correlated at +1, with substantial intra-, and inter-taxon variation (Figs. 8, 9; not all extreme values are visible in the boxplots). Mean co-occurrences of all taxa differed significantly between on-shelf and off-shelf (Wilcoxon Rank Sum test, all combinations p < 0.0001, except winter of 2019 at TR) and these on-shelf/off-shelf patterns differed between the NH and TR lines (Figs. 8, 9). On the NH line, across both seasons and years, correlation coefficients were more positive off-shelf compared to on-shelf, while on the TR line the pattern was more complex. Winter and summer 2018 correlations on the TR line were the opposite of the NH line, with more positive correlations on-shelf relative to off-shelf. In contrast, average TR on- and off-shelf correlations in winter 2019 were virtually indistinguishable, before transitioning in summer 2019 to a pattern similar to the NH line where off-shelf correlations were significantly higher than those on-shelf.

**Figure 8.**
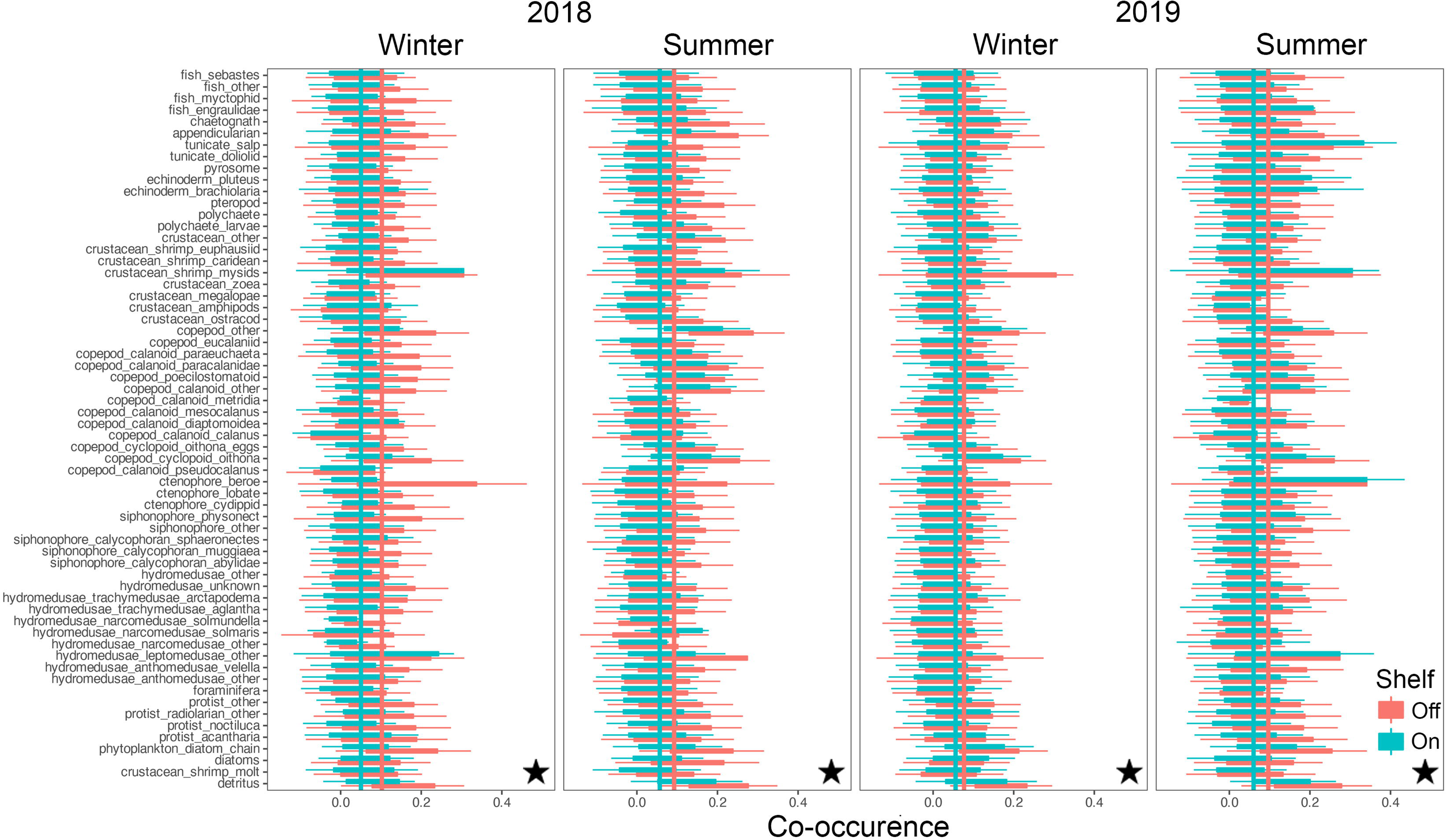
Co-occurrence of plankton taxa (y-axis) with all other taxa on the shelf (green) and off the shelf (red) along the Newport Hydrographic (NH) line. Vertical lines indicate taxa averages of co-occurrence as measured by spatial correlations. Stars indicate significant differences between on-, and off-shelf co-occurrence averages using Wilcoxon Rank Sum tests (p < 0.0001).

**Figure 9.**
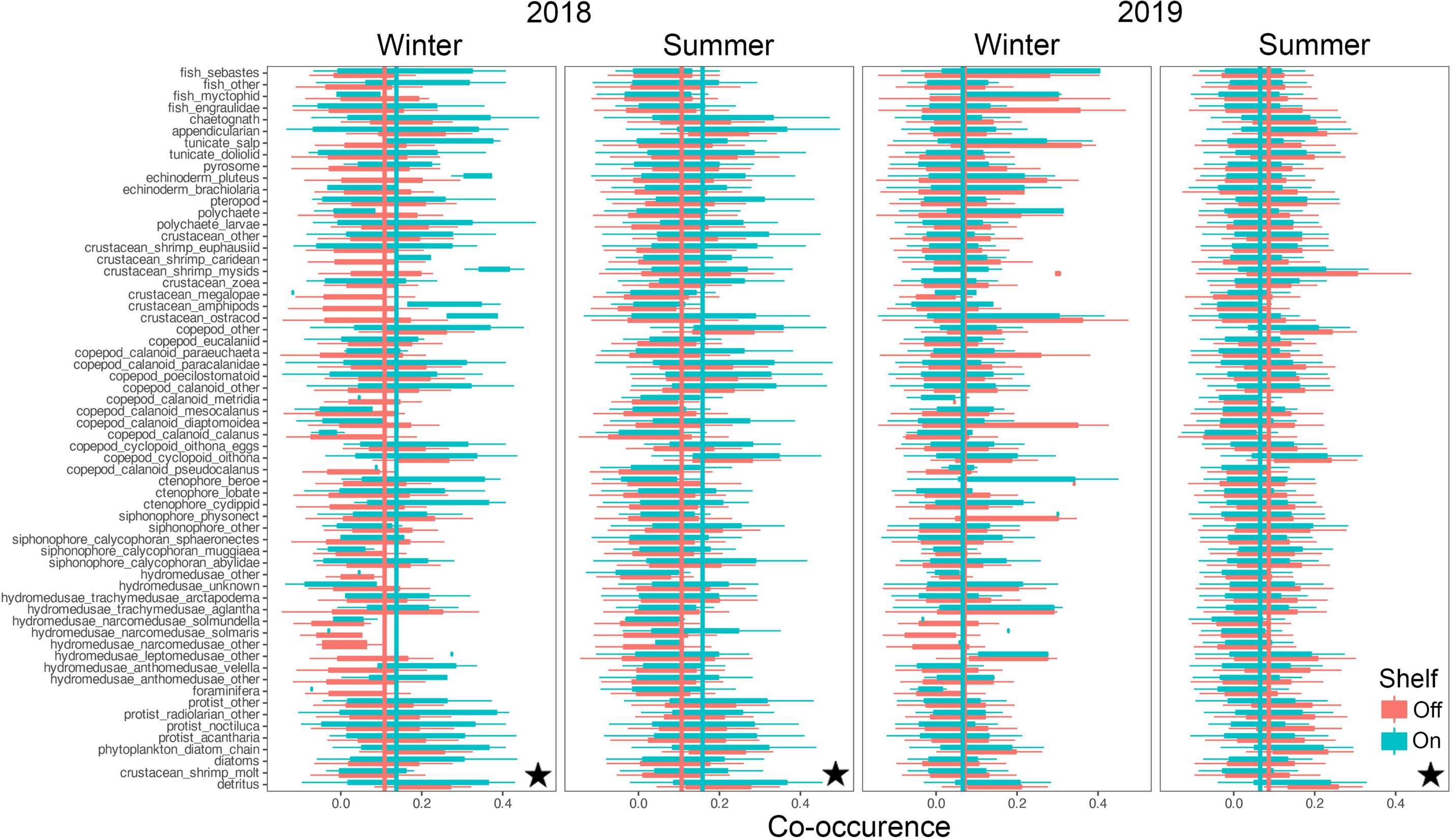
Co-occurrence of plankton taxa (y-axis) with all other taxa on the shelf (green) and off the shelf (red) along the Trinidad Head line. Vertical lines indicate taxa averages of co-occurrence as measured by spatial correlations. Stars indicate significant differences between on-, and off-shelf co-occurrence averages using Wilcoxon Rank Sum tests (p < 0.0001).

Similarly, the coefficients of variation (CVs) around these mean co-occurrences on the NH line were consistently higher on-shelf (> 2) compared to off-shelf (1.5), while on the TR line, the pattern was more complex (Fig. 10). In winter 2018, CVs for both on-shelf and off-shelf correlations on TR were virtually identical, while in summer 2018, the on-shelf CV was lower than off-shelf, before switching to a pattern similar to the NH line in 2019, with higher CVs on-shelf relative to off-shelf (Fig. 10). Mean co-occurrence of all taxa as a function of the preceding 10-d cumulative CUTI was significant on the TR Line (Fig. 11; Wilcoxon rank sum test p < 0.001), while on the NH line no such relationship was detected.

**Figure 10.**
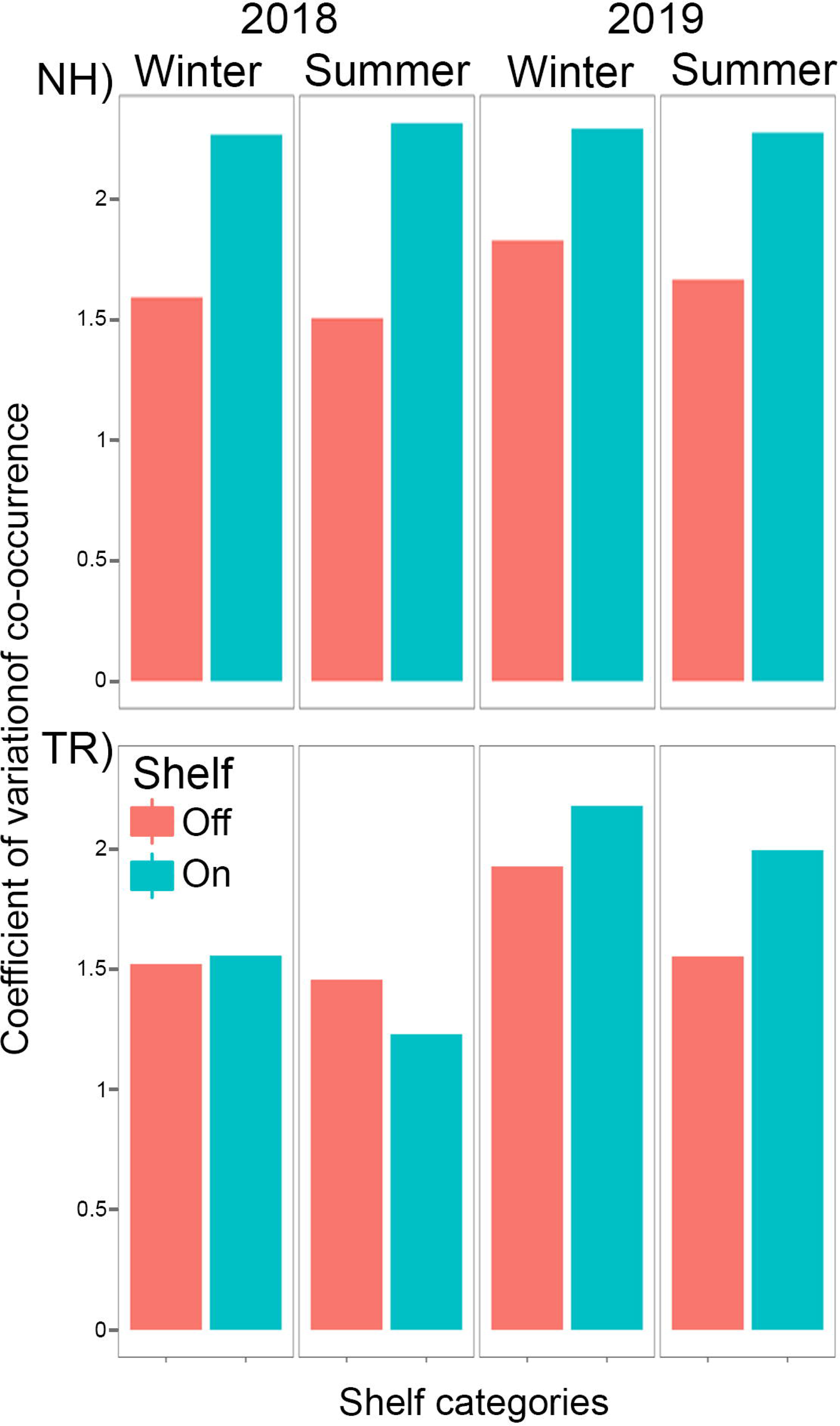
Coefficients of variation of plankton taxa co-occurrences along the Newport Hydrographic (NH) and Trinidad Head (TR) lines.

**Figure 11.**
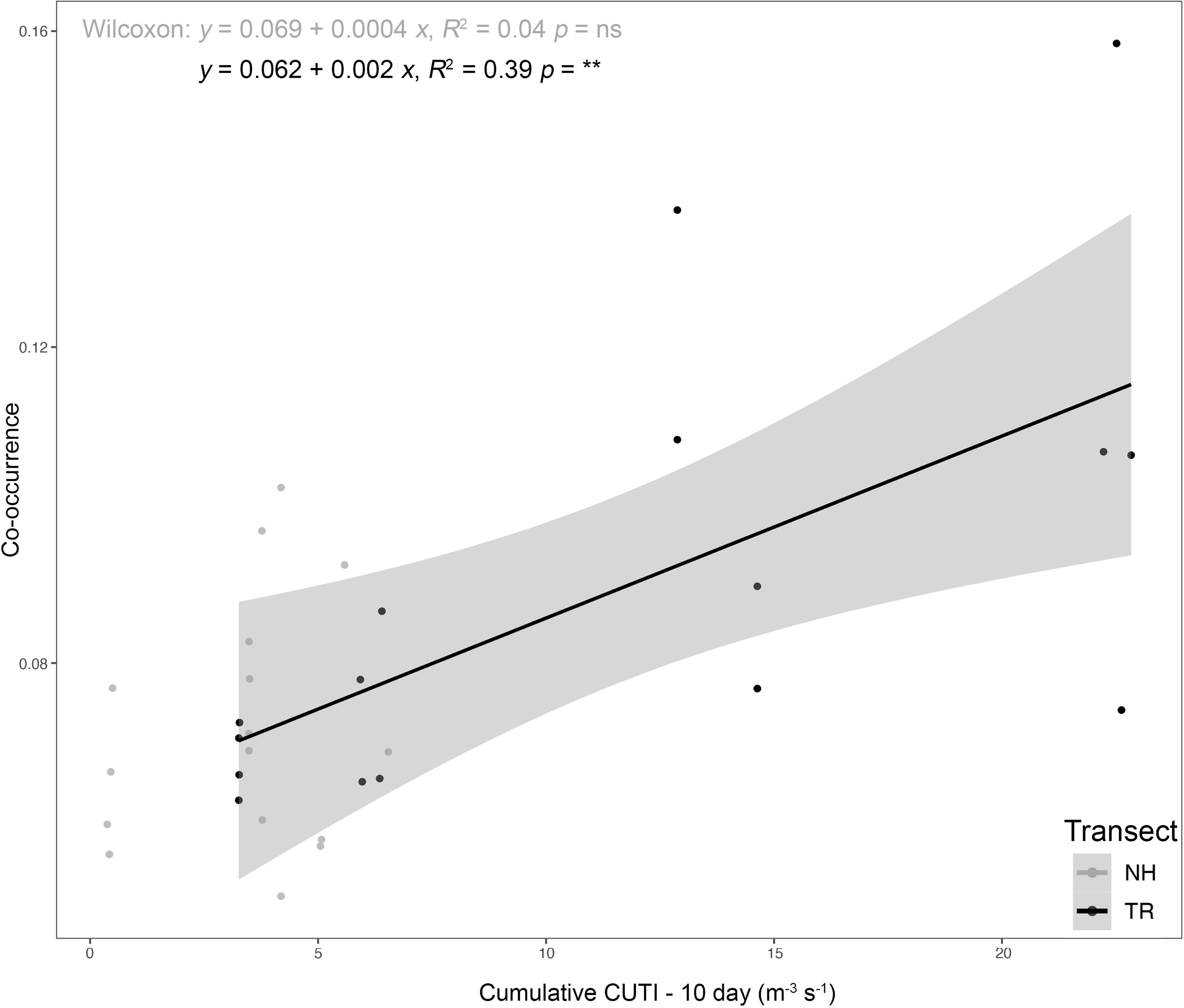
Plankton taxa co-occurrence as a function of the 10-d cumulative Coastal Upwelling Transport Index (CUTI) along the Newport Hydrographic (NH) and Trinidad Head (TR) lines; Wilcoxon test (ns = not significant, ** *p* < 0.001).

### 3.2 High spatial resolution taxa co-occurrence and their environmental drivers

The two Random Forests (RF) models designed to predict vertically and horizontally stratified taxa co-occurrences based on 83 biotic and abiotic variables (Table 1) explained 42% (NH) and 43% (TR) of the variance. The variables explaining most of the variance per model differed substantially between the two transect locations (Fig. 12). On the NH line, sampling depth was the most important predictor, followed by the binary on-shelf/off-shelf variable, temperature, density, the distance along the transect (i.e., how far offshore sampling occurred), and salinity (Fig. 12). These abiotic variables were followed by taxa concentrations of *Oithona* sp. copepods, appendicularians, other small copepods, and protists. On the TR line, the binary year variable (2018/2019) was the most important predictor, followed by the 10-d cumulative CUTI, the binary shelf indicator, sampling depth, the distance along the transect, oxygen, density, temperature, chl *a,* and salinity (Fig. 12).

**Figure 12.**
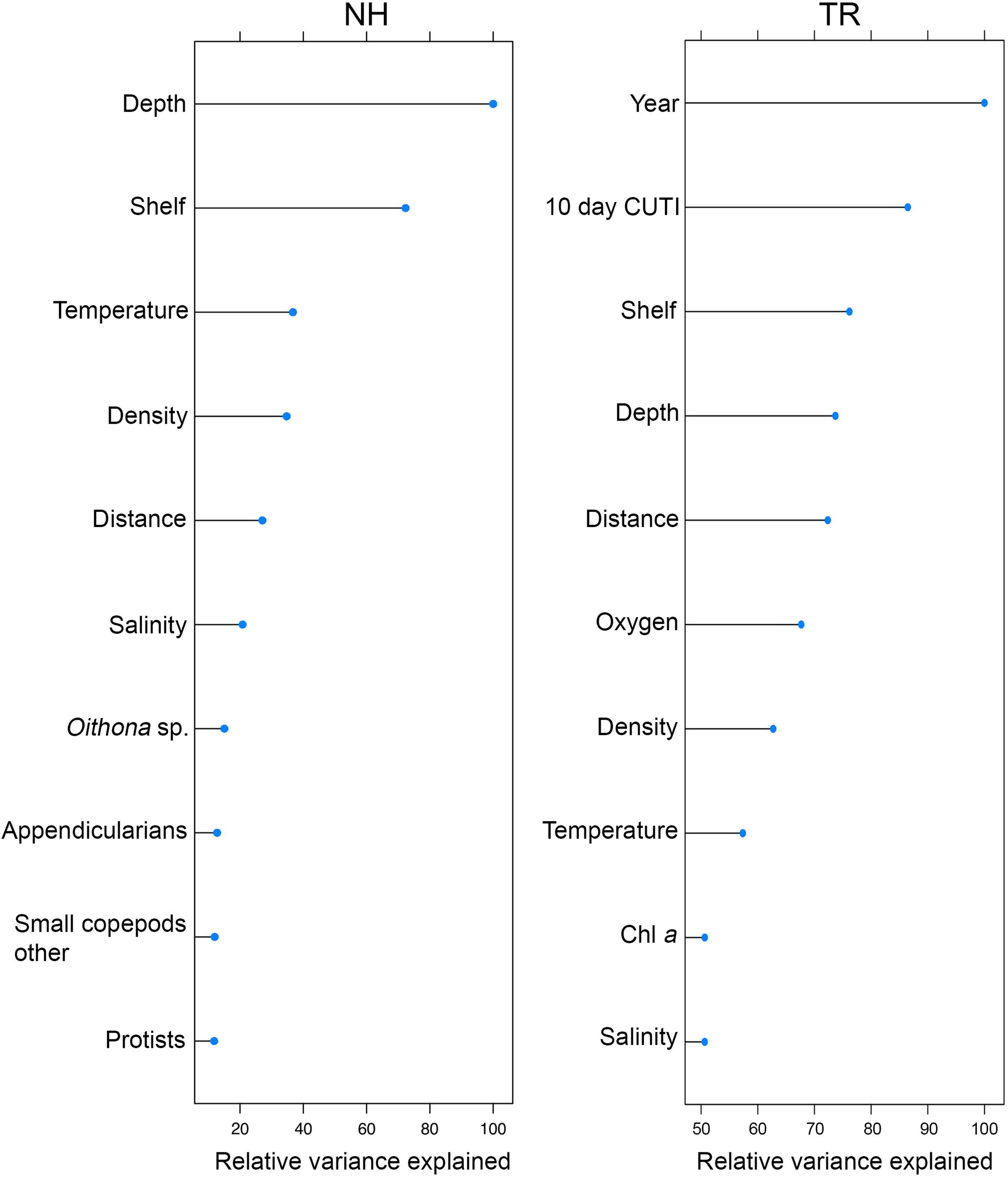
Top ten variables in the Random Forests models of plankton cooccurrence at the Newport Hydrographic (NH) and Trinidad Head (TR) lines, ordered by relative variance explained (variance scaled to 100% based on the most important variable).

On the NH line, a deeper sampling depth (> 85 m) led to a substantially higher chance of co-occurrence than in shallower water, while on-shelf in general predicted lower co-occurrences (Fig. 13). Temperature followed a gradual pattern where warmer temperatures predicted higher co-occurrence values. Density and salinity effects were similar in that the lowest densities and salinities led to the lowest co-occurrence and vice versa. Distance along the transect indicated that locations farther offshore predicted higher co-occurrences compared to inshore locations. Concentrations of *Oithona* copepods, appendicularians, other small copepods, and protists had similar effects in that their lowest concentrations predicted the least likely co-occurrence, followed by a rise in predicted co-occurrence as taxa concentrations increased. Protists showed the strongest such effect whereby a steady increase in protist concentrations led to the fastest increase in predicted co-occurrence, matched only by sampling depth.

**Figure 13.**
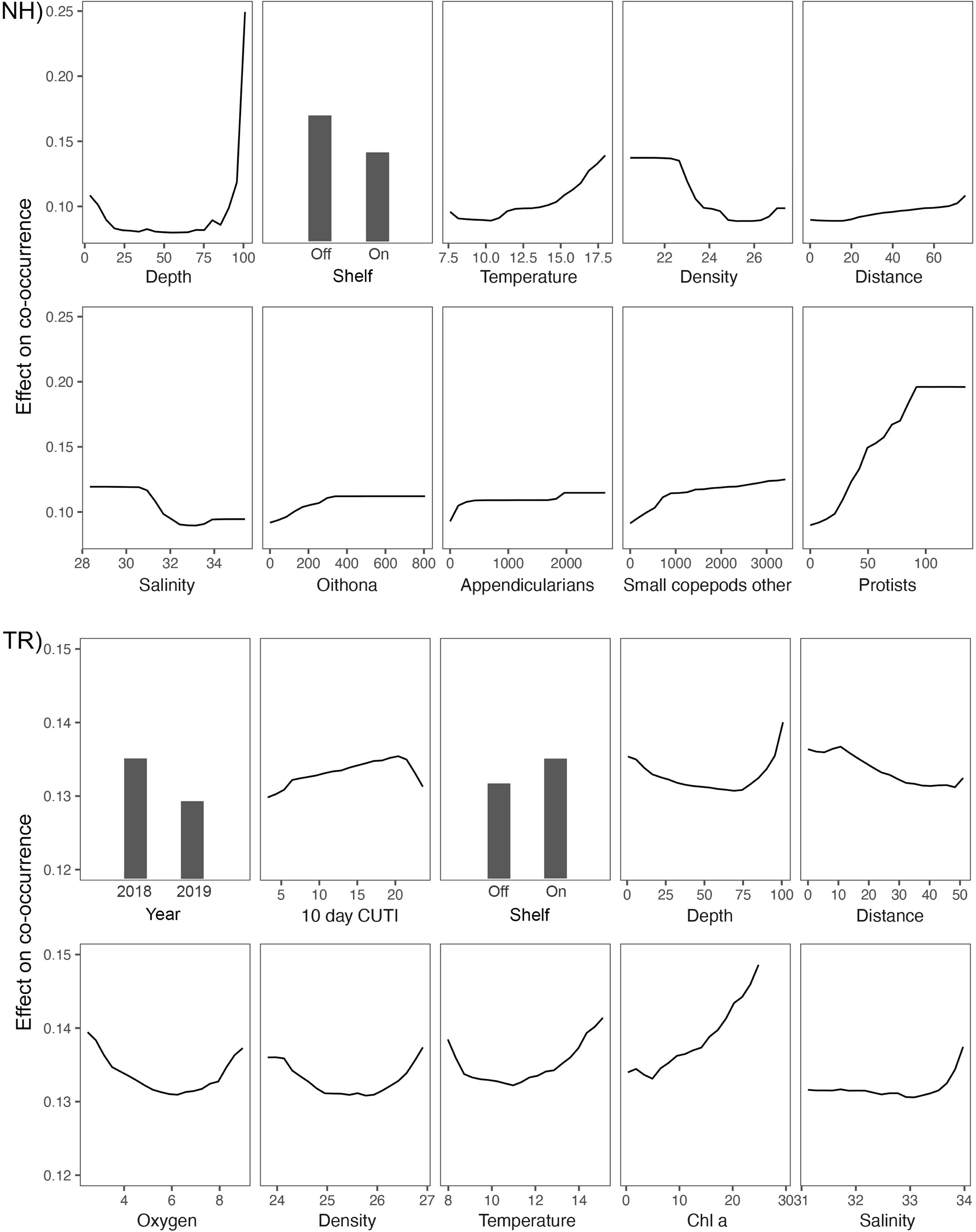
Partial dependence plots for the top 10 most important variables in the Random Forests models of plankton co-occurrence on the Newport Hydrographic (NH) and Trinidad Head (TR) lines. Note the differing y-axis scale between NH and TR

On the TR line, 2018 data were a good predictor of higher co-occurrence, while 2019 data led to lower values (Fig. 13). The 10-d cumulative CUTI was an important predictor and increasing CUTI values led to higher predicted co-occurrence until a CUTI of ∼20 m^-3^ s^-1^, after which the predicted co-occurrence dropped (Fig. 13). Notable differences between the TR and NH lines were that the shelf variable in the TR model showed that higher co-occurrence was predicted at on-shelf locations, which was followed by the distance variable that showed a decline in the predicted co-occurrence going from on-shore to off-shore. While the depth variable on the TR line also showed the highest predicted co-occurrence at 100m depth, the range of the predicted co-occurrences was substantially narrower than that on the NH line. Oxygen, density, and temperature partial effects plots had a very similar pattern between locations: the lowest and highest values generally led to the highest predicted co-occurrence. The positive effect of Chl *a* on the co-occurrence of taxa increased steadily across the spectrum of chl *a* values (>20 μg l^-1^). The salinity partial effects profile differed substantially between locations: in contrast to the NH line, the highest salinities on the TR line were good predictors of higher taxa co-occurrence.

## 4 Discussion

### 4.1 Plankton community structure in the Northern California Current

As the northern portion of the prototypical Eastern Boundary Upwelling System, the northern California Current (NCC) is characterized by strong, but intermittent upwelling, a typically short food web, and subsequently high fisheries biomass (Ryther, 1969; Pauly and Christensen, 1995; Rykaczewski and Checkley, 2008). While plankton community structure in the NCC has received much attention over the last decades (Peterson and Keister, 2003; Peterson et al., 2014, 2017; Brodeur et al., 2019; Weber et al., 2021; Thompson et al., 2022), we still lack a comprehensive understanding of how plankton community structure responds to changing environmental conditions. Our high-resolution imaging of the water column at two locations in the NCC that differ in their scale and continuity of upwelling enabled us to tease apart the relationships of new and recycled production (i.e., microbial loop) and plankton community structure. By simultaneously sampling a wide range of organisms including protists, phytoplankton, zooplankton, and fragile gelatinous plankton, in situ plankton imaging can bridge the sampling gap in studying the microbial and new production driven components of the plankton (Biard et al., 2016; Briseño-Avena et al., 2020; Schmid et al., 2020).

Using plankton co-occurrence as a proxy for community structure (Reese and Brodeur, 2006; Brodeur et al., 2008; Sildever et al., 2021; Costas-Selas et al., 2022), we condensed >2000 high resolution taxa distribution profiles into a unified community approach. Plankton co-occurrence patterns differed substantially between the two sampled locations, NH with intermittent upwelling and TR with more continuous upwelling. TR plankton co-occurrences in 2018 were higher on-shore relative to off-shore, consistent with the expectations for a nearshore upwelling system where new productivity is fueled by nutrients brought to the euphotic zone (Barth et al., 2007; Bograd et al., 2009; Jacox et al., 2018). However, TR plankton co-occurrences in 2019 differed from this pattern, likely induced by much lower upwelling and hence chl *a* in 2019. In sharp contrast to TR, plankton co-occurrence at NH was consistently higher in the more oligotrophic off-shelf waters (Peterson et al., 2017) relative to on-shelf waters. This pattern at NH remained consistent across all years and seasons, despite the lower upwelling in 2019 and suggests that in intermittent upwelling, nearshore conditions are generally less conducive to setting up a stable plankton community structure.

The relative importance of the microbial loop was highlighted by the high spatial resolution modeling of plankton co-occurrences. Among the variables explaining the most variance in taxa co-occurrence on the NH line over time were concentrations of several taxa associated with the microbial loop. *Oithona* sp copepods are small, ubiquitous cyclopoid copepods that are closely linked to the microbial loop through feeding on protozooplankton such as ciliates and dinoflagellates (Atienza et al, 2006; Zamora-Terol et al., 2014). Appendicularians similarly feed on the very small constituents of the microbial loop - down to picoplankton sizes - (Gorsky and Fenaux, 1998; Sutherland et al., 2010; Sutherland and Thompson, 2022) by using specialized feeding-filters (Conley and Sutherland, 2017). Appendicularians can be extremely abundant – we measured dense patches of >10,000 ind. m^-3^ on the TR line – and are important prey for numerous taxa, including copepods, chaetognaths, ctenophores, and larval to small adult fishes (Gorsky and Fenaux, 1998; Purcell et al., 2005). Being a key driver of plankton community structure on the NH line, while also accumulating in dense patches on the TR line, the presence of appendicularians indicates the constant underlying activity of the microbial loop. A key feature of appendicularians are their mucous houses that are discarded regularly and contribute significantly to vertical ocean carbon flux (Alldredge, 1976; Sato et al., 2003; Luo et al., 2022). The high importance of appendicularians in contributing to plankton community structure in intermittent upwelling systems further advances the body of literature emphasizing the often-overlooked importance of gelatinous plankton, and specifically, appendicularians. Protists are the prototypical constituent of the microbial loop (Azam et al., 1983; Williams and Ducklow, 2019; Glibert and Mitra, 2022) and their importance in generating plankton community structure at NH is not only a robust confirmation of high microbial loop activity but may also reflect the consumption of protists by appendicularians and *Oithona* copepods. These faunal patterns are consistent with the importance of sampling depth and the on-shelf/off-shelf variable in the NH co-occurrence model, as deeper off-shelf waters tend to be more oligotrophic and favorable for heightened microbial activity (Azam et al., 1983; Williams and Ducklow, 2019; Glibert and Mitra, 2022).

In contrast to the variables influencing plankton community structure at NH, variation in plankton co-occurrences at TR was influenced most strongly by upwelling and chl *a*, both indicative of a system dominated by new productivity with relatively reduced importance of the microbial loop, and generally shorter trophic pathways (Rykaczewski and Checkley, 2008; Jacox et al., 2018). While the positive effect of chl *a* on predicted plankton co-occurrence increased almost linearly across the range of observed chl *a* values, predicted co-occurrence increased with the cumulative 10-d CUTI only up to a value of ∼20 m^-3^ s^-1^ before dropping off. This non-linear relationship may be due to an imbalance of upwelling and relaxation events whereby too much and continuous upwelling lead to advective loss of plankton off the shelf (Largier et al., 2006; Kudela et al., 2008). Sampling year was also an important driver on the TR line where both upwelling strength and chl *a* were much higher in 2018 compared to 2019. Northern California upwelling and chl *a* levels in 2018 and 2019 have been reported as average and slightly below average, respectively (Thompson et al., 2018, 2019); however, our measurements reveal larger differences. At TR, cumulative CUTI was much higher in 2018 (winter = 13.8 m^-3^ s^-1^; summer = 22.3 m^-3^ s^-1^) relative to 2019 (winter = 3.3 m^-3^ s^-1^; summer = 6.2 m^-3^ s^-1^). This interannual difference in CUTI likely also led to the much higher chl *a* levels observed in 2018 relative to 2019 (> 45 μg l^-1^ in 2018 vs 7.5 μg l^-1^ in 2019). Considering that CUTI and chl *a* were both important predictors in the TR model, these large differences between 2018 and 2019 likely explain why the ‘year’ variable was also important and why 2018 predicted higher co-occurrences. Other variables that were important in driving plankton co-occurrences on the TR and NH lines were temperature and oxygen. Both are key drivers in structuring pelagic plankton ecosystems, through physiological effects that can impact predator-prey interactions, as well as physical discontinuities than can constrain plankton movement (Rutherford et al., 1999; Rebstock, 2003; Brodeur et al., 2019). Temperature and oxygen are also two of the variables most affected by climate change (Chan et al., 2019; Bograd et al., 2022; Smith et al., 2022).

In a strong (i.e., continuous) upwelling environment (TR), a reduction in upwelling and resulting lower chl *a* lead to a reversal of the prevailing on-shelf/off-shelf pattern of co-occurrence, while in an already lower upwelling strength environment (i.e., intermittent upwelling regime; NH), a further reduction of upwelling leads to little change in on-shelf/off-shelf co-occurrence patterns. The larger effect on plankton co-occurrences in the strong upwelling environment is consistent with the expectation that the established trophic web is reliant on the input of nutrients through upwelling and subsequent phytoplankton blooms (Barth et al., 2007; Bograd et al., 2009; Jacox et al., 2018). In an intermittent upwelling environment, where we found several microbial loop taxa to be important in predicting co-occurrences, the established trophic web (including a protist - *Oithona* - appendicularian link) is less reliant on nutrient input from upwelling (Azam et al., 1983; Williams and Ducklow, 2019), thus a further reduction in upwelling would be expected to have a smaller effect.

It is well established that the microbial loop is an important part of many marine ecosystems (Wilkerson et al., 1987; Taylor and Landry, 2018; Williams and Ducklow, 2019; Thompson et al., 2021; Glibert and Mitra, 2022). Recent establishment of the mixoplankton paradigm—ubiquitous microbes that survive on phototrophy and phagotrophy synergistically—has had far reaching ripple effects (Flynn et al., 2019; Glibert and Mitra, 2022). Long considered minor players, mixotrophs are now known to comprise large parts of the microbial loop and are of high importance in the global plankton trophic web. Nonetheless, the role of the microbial loop in shaping overall plankton community structure, particularly in the context of variable environmental conditions, is not well understood. Several comparative studies have investigated the relative carbon contributions of broad taxa to new productivity and the microbial loop (Tilstone et al., 1999; Vargas et al., 2007; Landry et al., 2012; Taylor et al., 2015). For example, in a productive coastal upwelling region in the Humboldt Current, the microbial loop was found to channel a large portion of the energy flow, while new productivity contributed only a small portion of the transferred carbon (Vargas et al., 2007).

Our in-situ plankton imagery demonstrates that the role of the microbial loop in driving mesoplankton community structure is more evident in intermittent upwelling regimes relative to continuous upwelling regions. While areas dominated by upwelling and high nutrient input also include microbial constituents, new productivity plays a larger role in structuring the plankton community. Here, large changes in upwelling result in sharp spatial changes to plankton community structure. In intermittent or low upwelling areas, microbial loop constituents are more important drivers of overall plankton community structure, resulting in a more temporally stable plankton community structure, even in the face of changes to upwelling strength.

Complexities of nutrient-plankton interactions, including the microbial loop, are often not well represented in models, and need refining, especially with regard to adequately including mixotrophy (Ratnarajah et al., 2023). Updating these models becomes especially urgent in the uncertain future ocean.

### 4.2 Plankton community structure under future climate change

Recently the NCC has been subject to disruptive marine heatwaves, affecting multiple trophic levels (Cavole et al., 2016; Oliver et al., 2018; Fennie et al., *in revision*) and reducing biodiversity on basin scales (Smale et al., 2019; Smith et al., 2022). Unfortunately, such extreme events are also predicted to become more prevalent in the future (Jacox et al., 2022). Marine heatwaves can lead to changes in plankton and nekton community structure (Brodeur et al., 2019) and to die-offs in seabirds, marine mammals, and kelp (Smith et al., 2022). Simultaneously, deeper and stronger stratification will result in lower nutrient supply to surface waters, with a resulting impact on food web structure—i.e., a shift to smaller plankters that rely to a greater extent on microbial-based nutrient recycling (Behrenfeld and Boss, 2013)—generating longer, less-efficient food chains. Meanwhile, changing wind patterns are projected to intensify upwelling in the NCC, and to decrease upwelling-favorable winds in the central and southern California Current Ecosystem (Buil et al., 2021). Our findings suggest that as these changes in wind patterns lead to shifts in intermittent and continuous upwelling regimes, current continuous upwelling regions will likely transition to a plankton community structure that is driven more by microbial loop constituents, and current intermittent upwelling regions will likely transition to systems dominated by new productivity. Such fundamental changes would likely have important consequences for energy transport through the trophic web to top predators and fisheries.

### 4.3 Conclusions

Collection and analysis of a vast dataset of in situ underwater plankton imagery (>1.1 billion plankton images) revealed substantial differences in the way that plankton community structure is driven under intermittent and continuous upwelling regimes. A reduction of upwelling strength in a continuous upwelling regime induced large scale changes in plankton community structure that affected on-shelf and off-shelf taxa co-occurrences, while in an intermittent upwelling regime, more strongly influenced by microbial loop constituents, a reduction of upwelling strength had little effect on plankton community structure. We thus hypothesize that high microbial loop activity enhances the resilience of plankton community structure to climate change induced shifts in upwelling strength. This concept is consistent with the mixotrophy paradigm in which the base of the microbial loop–the mixotrophs–are better adapted to a changing ocean (e.g., changing nutrient availability) than pure auto-, or heterotrophs, due to their ability to survive on either (Glibert and Mitra, 2022).

## 8 Author Contributions

MSS performed analyses and wrote the initial manuscript; SS, RKC, KRS, and MSS conceptualized hypotheses and research questions; SS, RKC, and KRS designed the study, sampling program, and wrote grant proposals; MSS, RKC, and KRS collected data; all authors interpreted the data, discussed the results, contributed to the critical revision of the manuscript and figures, and approved the final version.

## 10 Acknowledgments

We thank our collaborators Chris Sullivan and Dominic Daprano (both at Oregon State University) for invaluable support with processing the vast quantity of data presented here. We also thank Kelsey Swieca and Christian Briseño-Avena for their help creating the sCNN training library. Current and former members of the OSU Plankton Ecology Lab at the Hatfield Marine Science Center helped collect the data presented here on the four cruises, we thank them for their efforts – Jami Ivory, Christian Briseño-Avena, Kelsey Swieca, Miram Gleiber, H. Will Fennie, Megan Wilson, and Keely Axler. The professionalism of the officers and crews of RV *Sikuliaq* (Winter 2018 and 2019), RV *Sally Ride* (Summer 2018) and RV *Atlantis* (Summer 2019) enabled this project, and we are grateful to them. We thank NSF XSEDE/ACCESS and specifically staff Manu Shantharam at the San Diego Supercomputing Center as well as Sergiu Sanlievici at the Pittsburgh Supercomputing Center for their support with classifying the image data presented here.

## 12 Data Availability Statement

Data are available at NSF’s BCO-DMO https://www.bco-dmo.org/project/743417 as well as the R2R program: https://www.rvdata.us/search/cruise/SKQ201804S, https://www.rvdata.us/search/cruise/SR1810, https://www.rvdata.us/search/cruise/AT42-13, and https://www.rvdata.us/search/cruise/SKQ201903S.

## Footnotes

1 ; https://mjacox.com/upwelling-indices/

## 11 References

Alldredge, A. L. (1976). Discarded appendicularian houses as sources of food, surface habitats, and particulate organic matter in planktonic environments. Limnol Oceanogr 21, 14–24. doi: 10.4319/lo.1976.21.1.0014.

Atienza, D., Calbet, A., Saiz, E., Alcaraz, M., and Trepat, I. (2006). Trophic impact, metabolism, and biogeochemical role of the marine cladoceran *Penilia avirostris* and the co-dominant copepod *Oithona nana* in NW Mediterranean coastal waters. Mar Biol 150, 221–235. doi: 10.1007/s00227-006-0351-z.

Azam, F., Fenchel, T., Field, J., Gray, J., Meyer-Reil, L., and Thingstad, F. (1983). The ecological role of water-column microbes in the sea. Mar Ecol Prog Ser 10, 257–263. doi: 10.3354/meps010257.

Bakun, A. (1990). Global climate change and intensification of coastal ocean upwelling. Science 247, 198–201. doi: 10.1126/science.247.4939.198.

Bakun, A., Black, B. A., Bograd, S. J., García-Reyes, M., Miller, A. J., Rykaczewski, R. R., et al. (2015). Anticipated effects of climate change on coastal upwelling ecosystems. Curr Clim Change Reports 1, 85–93. doi: 10.1007/s40641-015-0008-4.

Bakun, A., and Nelson, C. S. (1991). The Seasonal Cycle of Wind-Stress Curl in Subtropical Eastern Boundary Current Regions. J Phys Oceanogr 21, 1815–1834. doi: 10.1175/1520-0485(1991)021<1815:tscows>2.0.co;2.

Barth, J. A., Menge, B. A., Lubchenco, J., Chan, F., Bane, J. M., Kirincich, A. R., et al. (2007). Delayed upwelling alters nearshore coastal ocean ecosystems in the northern California current. Proc National Acad Sci 104, 3719–3724. doi: 10.1073/pnas.0700462104.

Barth, J. A., Pierce, S. D., and Castelao, R. M. (2005). TimeLJdependent, windLJdriven flow over a shallow midshelf submarine bank. J Geophys Res Oceans 110. doi: 10.1029/2004jc002761.

Barth, J. A., Pierce, S. D., and Smith, R. L. (2000). A separating coastal upwelling jet at Cape Blanco, Oregon and its connection to the California Current System. Deep Sea Res Part Ii Top Stud Oceanogr 47, 783–810. doi: 10.1016/s0967-0645(99)00127-7.

Behrenfeld, M. J., and Boss, E. S. (2013). Resurrecting the Ecological Underpinnings of Ocean Plankton Blooms. Annu Rev Mar Sci 6, 167–194. doi: 10.1146/annurev-marine-052913-021325.

BenoitLJBird, K. J., Shroyer, E. L., and McManus, M. A. (2013). A critical scale in plankton aggregations across coastal ecosystems. Geophys Res Lett 40, 3968–3974. doi: 10.1002/grl.50747.

Biard, T., Stemmann, L., Picheral, M., Mayot, N., Vandromme, P., Hauss, H., et al. (2016). In situ imaging reveals the biomass of giant protists in the global ocean. Nature 532, 504. doi: 10.1038/nature17652

Bograd, S. J., Jacox, M. G., Hazen, E. L., Lovecchio, E., Montes, I., Buil, M. P., et al. (2022). Climate Change Impacts on Eastern Boundary Upwelling Systems. Annu Rev Mar Sci 15, 303–328. doi: 10.1146/annurev-marine-032122-021945.

Bograd, S. J., Schroeder, I., Sarkar, N., Qiu, X., Sydeman, W. J., and Schwing, F. B. (2009). Phenology of coastal upwelling in the California Current. Geophys Res Lett 36. doi: 10.1029/2008gl035933.

Breiman, L. (2001). Random Forests. Mach Learn 45, 5–32. doi: 10.1023/a:1010933404324.

Briseño-Avena, C., Schmid, M. S., Swieca, K., Sponaugle, S., Brodeur, R. D., and Cowen, R. K. (2020). Three-dimensional cross-shelf zooplankton distributions off the Central Oregon Coast during anomalous oceanographic conditions. Prog Oceanogr 188, 102436. doi: 10.1016/j.pocean.2020.102436.

Brodeur, R. D., Auth, T. D., and Phillips, A. J. (2019). Major Shifts in Pelagic Micronekton and Macrozooplankton Community Structure in an Upwelling Ecosystem Related to an Unprecedented Marine Heatwave. Frontiers Mar Sci 6, 212. doi: 10.3389/fmars.2019.00212.

Brodeur, R. D., Suchman, C. L., Reese, D. C., Miller, T. W., and Daly, E. A. (2008). Spatial overlap and trophic interactions between pelagic fish and large jellyfish in the northern California Current. Mar Biol 154, 649–659. doi: 10.1007/s00227-008-0958-3.

Brown, J. H., Gillooly, J. F., Allen, A. P., Savage, V. M., and West, G. B. (2004). Toward a metabolic theory of ecology. Ecology 85, 1771–1789. doi: 10.1890/03-9000.

Buil, M. P., Jacox, M. G., Fiechter, J., Alexander, M. A., Bograd, S. J., Curchitser, E. N., et al. (2021). A Dynamically Downscaled Ensemble of Future Projections for the California Current System. Frontiers Mar Sci 8, 612874. doi: 10.3389/fmars.2021.612874.

Cavan, E. L., Laurenceau-Cornec, E. C., Bressac, M., and Boyd, P. W. (2019). Exploring the ecology of the mesopelagic biological pump. Prog Oceanogr 176, 102125. doi: 10.1016/j.pocean.2019.102125.

Cavole, L., Demko, A., Diner, R., Giddings, A., Koester, I., et al. (2016). Biological Impacts of the 2013–2015 Warm-Water Anomaly in the Northeast Pacific: Winners, Losers, and the Future. Oceanography 29. doi: 10.5670/oceanog.2016.32.

Chan, F., Barth, J. A., Lubchenco, J., Kirincich, A., Weeks, H., Peterson, W. T., et al. (2008). Emergence of Anoxia in the California Current Large Marine Ecosystem. Science 319, 920–920. doi: 10.1126/science.1149016.

Chan, F., Barth, J., Kroeker, K., Lubchenco, J., and Menge, B. (2019). The Dynamics and Impact of Ocean Acidification and Hypoxia: Insights from Sustained Investigations in the Northern California Current Large Marine Ecosystem. Oceanography 32, 62–71. doi: 10.5670/oceanog.2019.312.

Checkley, D. M., and Barth, J. A. (2009). Patterns and processes in the California Current System. Prog Oceanogr 83, 49–64. doi: 10.1016/j.pocean.2009.07.028.

Chesson, P. (2000). Mechanisms of maintenance of species diversity. Annu Rev Ecol Syst 31, 343–366. doi: 10.1146/annurev.ecolsys.31.1.343.

Conley, K. R., and Sutherland, K. R. (2017). Particle shape impacts export and fate in the ocean through interactions with the globally abundant appendicularian Oikopleura dioica. Plos One 12, e0183105. doi: 10.1371/journal.pone.0183105.

Costas-Selas, C., Martínez-García, S., Logares, R., Hernández-Ruiz, M., and Teira, E. (2022). Role of Bacterial Community Composition as a Driver of the Small-Sized Phytoplankton Community Structure in a Productive Coastal System. Microbial Ecol, 1–18. doi:10.1007/s00248-022-02125-2.

Cowen, R. K., Greer, A. T., Guigand, C. M., Hare, J. A., Richardson, D. E., and Walsh, H. J. (2013). Evaluation of the In Situ Ichthyoplankton Imaging System (ISIIS): comparison with the traditional (bongo net) sampler. Fish B 111. doi: 10.7755/fb.111.1.1.

Cowen, R. K., and Guigand, C. M. (2008). In situ ichthyoplankton imaging system (ISIIS): system design and preliminary results. Limnology Oceanogr Methods 6, 126–132. doi: 10.4319/lom.2008.6.126.

Denman, K. L., and Gargett, A. E. (1995). Biological-Physical Interactions in the Upper Ocean: The Role of Vertical and Small Scale Transport Processes. Annu Rev Fluid Mech 27, 225–256. doi: 10.1146/annurev.fl.27.010195.001301.

Dickey, T. D., and Bidigare, R. R. (2005). Interdisciplinary oceanographic observations: the wave of the future. Sci Mar 69, 23–42. doi: 10.3989/scimar.2005.69s123.

Dickson, M., and Wheeler, P. (1995). Ammonium uptake and regeneration rates in a coastal upwelling regime. Mar Ecol Prog Ser 121, 239–248. doi: 10.3354/meps121239.

Doney, S. C., Busch, D. S., Cooley, S. R., and Kroeker, K. J. (2020). The Impacts of Ocean Acidification on Marine Ecosystems and Reliant Human Communities. Annu Rev Env Resour 45, 1–30. doi: 10.1146/annurev-environ-012320-083019.

Doney, S. C., Ruckelshaus, M., Duffy, J. E., Barry, J. P., Chan, F., English, C. A., et al. (2012). Climate Change Impacts on Marine Ecosystems. Annu Rev Mar Sci 4, 11–37. doi: 10.1146/annurev-marine-041911-111611.

Faillettaz, R., Picheral, M., Luo, J. Y., Guigand, C., Cowen, R. K., and Irisson, J.-O. (2016). Imperfect automatic image classification successfully describes plankton distribution patterns. Methods Oceanogr 15, 60–77. doi: 10.1016/j.mio.2016.04.003.

Feely, R. A., Sabine, C. L., Hernandez-Ayon, J. M., Ianson, D., and Hales, B. (2008). Evidence for Upwelling of Corrosive “Acidified” Water onto the Continental Shelf. Science 320, 1490–1492. doi: 10.1126/science.1155676.

Feinberg, L. R., and Peterson, W. T. (2003). Variability in duration and intensity of euphausiid spawning off central Oregon, 1996–2001. Prog Oceanogr 57, 363–379. doi: 10.1016/s0079-6611(03)00106-x.

Fennie, H.W., Grorud-Colvert, K., and Sponaugle, S. (*in revision*). Larval rockfish growth and survival in response to anomalous ocean conditions. Sci Rep

Flynn, K. J., Mitra, A., Anestis, K., Anschütz, A. A., Calbet, A., Ferreira, G. D., et al. (2019). Mixotrophic protists and a new paradigm for marine ecology: where does plankton research go now? J Plankton Res 41, 375–391. doi: 10.1093/plankt/fbz026.

GarcíaLJReyes, M., and Largier, J. L. (2012). Seasonality of coastal upwelling off central and northern California: New insights, including temporal and spatial variability. J Geophys Res Oceans 117, n/a-n/a. doi: 10.1029/2011jc007629.

Glibert, P. M., and Mitra, A. (2022). From webs, loops, shunts, and pumps to microbial multitasking: Evolving concepts of marine microbial ecology, the mixoplankton paradigm, and implications for a future ocean. Limnol Oceanogr 67, 585–597. doi: 10.1002/lno.12018.

González, H. E., Giesecke, R., Vargas, C. A., Pavez, M., Iriarte, J., Santibáñez, P., et al. (2004). Carbon cycling through the pelagic foodweb in the northern Humboldt Current off Chile (23°S). Ices J Mar Sci 61, 572–584. doi: 10.1016/j.icesjms.2004.03.021.

Gorsky, G., and Fenaux, R. (1998). “The role of Appendicularia in marine food webs,” in The Biology of Pelagic Tunicates (Oxford: Oxford University Press), 161–169.

Graham, B. (2014). Fractional Max-Pooling. Arxiv. doi: 10.48550/arxiv.1412.6071.

Greer, A. T., Chiaverano, L. M., Treible, L. M., Briseño-Avena, C., and Hernandez, F. J. (2021). From spatial pattern to ecological process through imaging zooplankton interactions. Ices J Mar Sci 78, 2664–2674. doi: 10.1093/icesjms/fsab149.

Greer, A. T., Schmid, M. S., Duffy, P. I., Robinson, K. L., Genung, M. A., Luo, J. Y., et al. (2023). In situ imaging across ecosystems to resolve the fineLJscale oceanographic drivers of a globally significant planktonic grazer. Limnol Oceanogr 68, 192–207. doi: 10.1002/lno.12259.

Hales, B., KarpLJBoss, L., Perlin, A., and Wheeler, P. A. (2006). Oxygen production and carbon sequestration in an upwelling coastal margin. Global Biogeochem Cy 20. doi: 10.1029/2005gb002517.

Haury, L. R., McGowan, J. A., and Wiebe, P. H. (1978). Spatial Pattern in Plankton Communities. 277–327. doi: 10.1007/978-1-4899-2195-6_12.

Hickey, B., and Banas, N. (2008). Why is the Northern End of the California Current System So Productive? Oceanography 21, 90–107. doi: 10.5670/oceanog.2008.07.

Hickey, B. M., and Banas, N. S. (2003). Oceanography of the U.S. Pacific Northwest Coastal Ocean and estuaries with application to coastal ecology. Estuaries 26, 1010–1031. doi: 10.1007/bf02803360.

HilleRisLambers, J., Adler, P. B., Harpole, W. S., Levine, J. M., and Mayfield, M. M. (2012). Rethinking Community Assembly through the Lens of Coexistence Theory. Ecol Evol Syst 43, 227–248. doi: 10.1146/annurev-ecolsys-110411-160411.

Irisson, J.-O., Ayata, S.-D., Lindsay, D. J., Karp-Boss, L., and Stemmann, L. (2021). Machine Learning for the Study of Plankton and Marine Snow from Images. Annu Rev Mar Sci 14, 1–25. doi: 10.1146/annurev-marine-041921-013023.

Jacox, M. G., Alexander, M. A., Amaya, D., Becker, E., Bograd, S. J., Brodie, S., et al. (2022). Global seasonal forecasts of marine heatwaves. Nature 604, 486–490. doi: 10.1038/s41586-022-04573-9.

Jacox, M. G., Edwards, C. A., Hazen, E. L., and Bograd, S. J. (2018). Coastal Upwelling Revisited: Ekman, Bakun, and Improved Upwelling Indices for the U.S. West Coast. J Geophys Res Oceans 123, 7332–7350. doi: 10.1029/2018jc014187.

Kara, A. B., Rochford, P. A., and Hurlburt, H. E. (2000). An optimal definition for ocean mixed layer depth. J Geophys Res Oceans 105, 16803–16821. doi: 10.1029/2000jc900072.

Kirchman, D. L. (2010). Microbial Ecology of the Oceans. 1–26. doi: 10.1002/9780470281840.ch1.

Kirincich, A. R., Barth, J. A., Grantham, B. A., Menge, B. A., and Lubchenco, J. (2005). WindLJdriven innerLJshelf circulation off central Oregon during summer. J Geophys Res Oceans 110. doi: 10.1029/2004jc002611.

Kudela, R., Banas, N., Barth, J., Frame, E., Jay, D., Largier, J., et al. (2008). New Insights into the Controls and Mechanisms of Plankton Productivity in Coastal Upwelling Waters of the Northern California Current System. Oceanography 21, 46–59. doi: 10.5670/oceanog.2008.04.

Kuhn, M. (2008). Building Predictive Models in R Using the caret Package. J Stat Softw 28. doi: 10.18637/jss.v028.i05.

Landry, M. R., Ohman, M. D., Goericke, R., Stukel, M. R., Barbeau, K. A., Bundy, R., et al. (2012). Pelagic community responses to a deep-water front in the California Current Ecosystem: overview of the A-Front Study. J Plankton Res 34, 739–748. doi: 10.1093/plankt/fbs025.

Largier, J. L., Lawrence, C. A., Roughan, M., Kaplan, D. M., Dever, E. P., Dorman, C. E., et al. (2006). WEST: A northern California study of the role of wind-driven transport in the productivity of coastal plankton communities. Deep Sea Res Part Ii Top Stud Oceanogr 53, 2833–2849. doi: 10.1016/j.dsr2.2006.08.018.

Lentz, S. J., and Chapman, D. C. (1989). Seasonal differences in the current and temperature variability over the northern California Shelf during the Coastal Ocean Dynamics Experiment. J Geophys Res Oceans 94, 12571–12592. doi: 10.1029/jc094ic09p12571.

Lertvilai, P., and Jaffe, J. S. (2022). In situ size and motility measurement of aquatic invertebrates with an underwater stereoscopic camera system using tilted lenses. Methods Ecol Evol 13, 1192– 1200. doi: 10.1111/2041-210x.13855.

Lima-Mendez, G., Faust, K., Henry, N., Decelle, J., Colin, S., Carcillo, F., et al. (2015). Determinants of community structure in the global plankton interactome. Science 348, 1262073. doi: 10.1126/science.1262073.

Lindegren, M., Thomas, M. K., Jónasdóttir, S. H., Nielsen, T. G., and Munk, P. (2020). Environmental niche separation promotes coexistence among ecologically similar zooplankton species—North Sea copepods as a case study. Limnol Oceanogr 65, 545–556. doi: 10.1002/lno.11322.

Lombard, F., Boss, E., Waite, A. M., Vogt, M., Uitz, J., Stemmann, L., et al. (2019). Globally Consistent Quantitative Observations of Planktonic Ecosystems. Frontiers Mar Sci 6, 196. doi: 10.3389/fmars.2019.00196.

Luo, J., Grassian, B., Tang, D., Irisson, J., Greer, A., Guigand, C., et al. (2014). Environmental drivers of the fine-scale distribution of a gelatinous zooplankton community across a mesoscale front. Mar Ecol Prog Ser 510, 129–149. doi: 10.3354/meps10908.

Luo, J. Y., Irisson, J., Graham, B., Guigand, C., Sarafraz, A., Mader, C., et al. (2018). Automated plankton image analysis using convolutional neural networks. Limnology Oceanogr Methods 16, 814–827. doi: 10.1002/lom3.10285.

Luo, J. Y., Stock, C. A., Henschke, N., Dunne, J. P., and O’Brien, T. D. (2022). Global ecological and biogeochemical impacts of pelagic tunicates. Prog Oceanogr 205, 102822. doi: 10.1016/j.pocean.2022.102822.

MacArthur, R. H. (1958). Population Ecology of Some Warblers of Northeastern Coniferous Forests. Ecology 39, 599–619. doi: 10.2307/1931600.

Mantua, N. J., and Hare, S. R. (2002). The Pacific Decadal Oscillation. J Oceanogr 58, 35–44. doi: 10.1023/a:1015820616384.

McClatchie, S., Cowen, R., Nieto, K., Greer, A., Luo, J. Y., Guigand, C., et al. (2012). Resolution of fine biological structure including small narcomedusae across a front in the Southern California Bight. J Geophys Res Oceans 117, n/a-n/a. doi: 10.1029/2011jc007565.

Mousseau, L., Fortier, L., and Legendre, L. (1998). Annual production of fish larvae and their prey in relation to size-fractionated primary production (Scotian Shelf, NW Atlantic). Ices J Mar Sci 55, 44–57. doi: 10.1006/jmsc.1997.0224.

Nagelkerken, I., and Connell, S. D. (2022). Ocean acidification drives global reshuffling of ecological communities. Global Change Biol 28, 7038–7048. doi: 10.1111/gcb.16410.

Ohman, M. D., Davis, R. E., Sherman, J. T., Grindley, K. R., Whitmore, B. M., Nickels, C. F., et al. (2019). Zooglider: An autonomous vehicle for optical and acoustic sensing of zooplankton. Limnology Oceanogr Methods 17, 69–86. doi: 10.1002/lom3.10301.

Oliver, E. C. J., Donat, M. G., Burrows, M. T., Moore, P. J., Smale, D. A., Alexander, L. V., et al. (2018). Longer and more frequent marine heatwaves over the past century. Nat Commun 9, 1324. doi: 10.1038/s41467-018-03732-9.

Orenstein, E. C., Ratelle, D., BriseñoLJAvena, C., Carter, M. L., Franks, P. J. S., Jaffe, J. S., et al. (2020). The Scripps Plankton Camera system: A framework and platform for in situ microscopy. Limnology Oceanogr Methods 18, 681–695. doi: 10.1002/lom3.10394.

Ortner, P. B., Cummings, S. R., Aftring, R. P., and Edgerton, H. E. (1979). Silhouette photography of oceanic zooplankton. Nature 277, 50–51. doi: 10.1038/277050a0.

Pauly, D., and Christensen, V. (1995). Primary production required to sustain global fisheries. Nature 374, 255–257. doi: 10.1038/374255a0.

Peterson, W., Fisher, J., Peterson, J., Morgan, C., Burke, B., et al. (2014). Applied Fisheries Oceanography: Ecosystem Indicators of Ocean Conditions Inform Fisheries Management in the California Current. Oceanography 27, 80–89. doi: 10.5670/oceanog.2014.88.

Peterson, W. T., Fisher, J. L., Strub, P. T., Du, X., Risien, C., Peterson, J., et al. (2017). The pelagic ecosystem in the Northern California Current off Oregon during the 2014–2016 warm anomalies within the context of the past 20 years. J Geophys Res Oceans 122, 7267–7290. doi: 10.1002/2017jc012952.

Peterson, W. T., and Keister, J. E. (2003). Interannual variability in copepod community composition at a coastal station in the northern California Current: a multivariate approach. Deep Sea Res Part Ii Top Stud Oceanogr 50, 2499–2517. doi: 10.1016/s0967-0645(03)00130-9.

Peterson, W. T., and Miller, C. B. (1975). Year-to-year variations in the planktology of the Oregon upwelling zone. Fish Bull 73, 642–653.

Picheral, M., Catalano, C., Brousseau, D., Claustre, H., Coppola, L., Leymarie, E., et al. (2022). The Underwater Vision Profiler 6: an imaging sensor of particle size spectra and plankton, for autonomous and cabled platforms. Limnology Oceanogr Methods 20, 115–129. doi: 10.1002/lom3.10475.

Pomeroy, L. R. (1974). The Ocean’s Food Web, A Changing Paradigm. Bioscience 24, 499–504. doi: 10.2307/1296885.

Prairie, J. C., Sutherland, K. R., Nickols, K. J., and Kaltenberg, A. M. (2012). Biophysical interactions in the plankton: A crossLJscale review. Limnology Oceanogr Fluids Environ 2, 121–145. doi: 10.1215/21573689-1964713.

Purcell, J.E., Sturdevant, M.V., Galt, C.P. (2005). A review of appendicularians as prey of fish and invertebrate predators, in: Gorsky, G., Youngbluth, M.J., Diebel, D. (Eds.), in Response of Marine Ecosystems to Global Change: Ecological Impact of Appendicularians. Contemporary Publishing International, Paris, France, pp. 359–435.

Ratnarajah, L., Abu-Alhaija, R., Atkinson, A., Batten, S., Bax, N. J., Bernard, K. S., et al. (2023). Monitoring and modelling marine zooplankton in a changing climate. Nat Commun 14, 564. doi: 10.1038/s41467-023-36241-5.

Rebstock, G. A. (2003). Long-term change and stability in the California Current System: lessons from CalCOFI and other long-term data sets. Deep Sea Res Part Ii Top Stud Oceanogr 50, 2583–2594. doi: 10.1016/s0967-0645(03)00124-3.

Reese, D. C., and Brodeur, R. D. (2006). Identifying and characterizing biological hotspots in the northern California Current. Deep Sea Res Part Ii Top Stud Oceanogr 53, 291–314. doi: 10.1016/j.dsr2.2006.01.014.

Ríos-Castro, R., Costas-Selas, C., Pallavicini, A., Vezzulli, L., Novoa, B., Teira, E., et al. (2022). Co-occurrence and diversity patterns of benthonic and planktonic communities in a shallow marine ecosystem. Frontiers Mar Sci 9, 934976. doi: 10.3389/fmars.2022.934976.

Robertson, R. R., and Bjorkstedt, E. P. (2020). Climate-driven variability in Euphausia pacifica size distributions off northern California. Prog Oceanogr 188, 102412. doi: 10.1016/j.pocean.2020.102412.

Robinson, K. L., Sponaugle, S., Luo, J. Y., Gleiber, M. R., and Cowen, R. K. (2021). Big or small, patchy all: Resolution of marine plankton patch structure at micro-to submesoscales for 36 taxa. Sci Adv 7, eabk2904. doi: 10.1126/sciadv.abk2904.

Rutherford, S., D’Hondt, S., and Prell, W. (1999). Environmental controls on the geographic distribution of zooplankton diversity. Nature 400, 749–753. doi: 10.1038/23449.

Rykaczewski, R. R., and Checkley, D. M. (2008). Influence of ocean winds on the pelagic ecosystem in upwelling regions. Proc National Acad Sci 105, 1965–1970. doi: 10.1073/pnas.0711777105.

Ryther, J. H. (1969). Photosynthesis and Fish Production in the Sea. Science 166, 72–76. doi: 10.1126/science.166.3901.72.

Sato, R., Tanaka, Y., and Ishimaru, T. (2003). Species-specific house productivity of appendicularians. Mar Ecol Prog Ser 259, 163–172. doi: 10.3354/meps259163.

Schmid, M. S., Cowen, R. K., Robinson, K., Luo, J. Y., Briseño-Avena, C., and Sponaugle, S. (2020). Prey and predator overlap at the edge of a mesoscale eddy: fine-scale, in-situ distributions to inform our understanding of oceanographic processes. Sci Rep 10, 921. doi: 10.1038/s41598-020-57879-x.

Schmid, M. S., Daprano, D., Jacobson, K. M., Sullivan, C., Briseño-Avena, C., Luo, J. Y., et al. (2021). A Convolutional Neural Network based high-throughput image classification pipeline - code and documentation to process plankton underwater imagery using local HPC infrastructure and NSF’s XSEDE. Zenodo. doi: http://dx.doi.org/10.5281/zenodo.4641158.

Schmid, M. S., and Fortier, L. (2019). The intriguing co-distribution of the copepods Calanus hyperboreus and Calanus glacialis in the subsurface chlorophyll maximum of Arctic seas. Elem Sci Anth 7, 50. doi: 10.1525/elementa.388.

Schmid, M. S., Maps, F., and Fortier, L. (2018). Lipid load triggers migration to diapause in Arctic Calanus copepods—insights from underwater imaging. J Plankton Res 40, 311–325. doi: 10.1093/plankt/fby012.

Shaw, C. T., Peterson, W. T., and Feinberg, L. R. (2010). Growth of Euphausia pacifica in the upwelling zone off the Oregon coast. Deep Sea Res Part Ii Top Stud Oceanogr 57, 584–593. doi: 10.1016/j.dsr2.2009.10.008.

Sildever, S., Laas, P., Kolesova, N., Lips, I., Lips, U., and Nagai, S. (2021). Plankton biodiversity and species co-occurrence based on environmental DNA – a multiple marker study. Metabarcoding Metagenomics 5, e72371. doi: 10.3897/mbmg.5.72371.

Smale, D. A., Wernberg, T., Oliver, E. C. J., Thomsen, M., Harvey, B. P., Straub, S. C., et al. (2019). Marine heatwaves threaten global biodiversity and the provision of ecosystem services. Nat Clim Change 9, 306–312. doi: 10.1038/s41558-019-0412-1.

Smetacek, V. (2012). Making sense of ocean biota: How evolution and biodiversity of land organisms differ from that of the plankton. J Biosciences 37, 589–607. doi: 10.1007/s12038-012-9240-4.

Smith, K. E., Burrows, M. T., Hobday, A. J., King, N. G., Moore, P. J., Gupta, A. S., et al. (2022). Biological Impacts of Marine Heatwaves. Annu Rev Mar Sci 15, 119–145. doi: 10.1146/annurev-marine-032122-121437.

Song, J., Bi, H., Cai, Z., Cheng, X., He, Y., Benfield, M. C., et al. (2020). Early warning of Noctiluca scintillans blooms using in-situ plankton imaging system: An example from Dapeng Bay, P.R. China. Ecol Indic 112, 106123. doi: 10.1016/j.ecolind.2020.106123.

Spitz, Y. H., Allen, J. S., and Gan, J. (2005). Modeling of ecosystem processes on the Oregon shelf during the 2001 summer upwelling. J Geophys Res Oceans 110. doi: 10.1029/2005jc002870.

Sutherland, K. R., Madin, L. P., and Stocker, R. (2010). Filtration of submicrometer particles by pelagic tunicates. Proc National Acad Sci 107, 15129–15134. doi: 10.1073/pnas.1003599107.

Sutherland, K. R., and Thompson, A. W. (2022). Pelagic tunicate grazing on marine microbes revealed by integrative approaches. Limnol Oceanogr 67, 102–121. doi: 10.1002/lno.11979.

Swieca, K., Sponaugle, S., Briseño-Avena, C., Schmid, M., Brodeur, R., and Cowen, R. (2020). Changing with the tides: fine-scale larval fish prey availability and predation pressure near a tidally modulated river plume. Mar Ecol Prog Ser 650, 217–238. doi: 10.3354/meps13367.

Swieca, K., Sponaugle, S., Schmid, M. S., Ivory, J., and Cowen, R. K. (in review). Shedding light on lanternfish: growth and diet of a larval myctophid across distinct upwelling regimes in the California Current. ICES Journal of Marine Science.

Taylor, A., and Landry, M. (2018). Phytoplankton biomass and size structure across trophic gradients in the southern California Current and adjacent ocean ecosystems. Mar Ecol Prog Ser 592, 1–17. doi: 10.3354/meps12526.

Taylor, A. G., Landry, M. R., Selph, K. E., and Wokuluk, J. J. (2015). Temporal and spatial patterns of microbial community biomass and composition in the Southern California Current Ecosystem. Deep Sea Res Part II Top Stud Oceanogr 112, 117–128. doi: 10.1016/j.dsr2.2014.02.006.

Thompson, A. R., Bjorkstedt, E. P., Bograd, S. J., Fisher, J. L., Hazen, E. L., Leising, A., et al. (2022). State of the California Current Ecosystem in 2021: Winter is coming? Frontiers Mar Sci 9, 958727. doi: 10.3389/fmars.2022.958727.

Thompson, A. R., Schroeder, I. D., Bograd, S. J., Hazen, E. L., Jacox, M. G., Leising, A., et al. (2019). State of the California current 2018–19: a novel anchovy regime and a new marine heat wave? Calif. Cooperat. Ocean. Fish. Investig. Rep. 60, 1–65.

Thompson, A. R., Schroeder, I. D., Bograd, S. J., Hazen, E. L., Jacox, M. G., Leising, A., et al. (2018). State of the California Current 2017-18: Still not quite normal in the North and getting interesting in the South. Calif. Cooperat. Ocean. Fish. Investig. Rep. 59, 1–66.

Thompson, A. W., Ward, A. C., Sweeney, C. P., and Sutherland, K. R. (2021). Host-specific symbioses and the microbial prey of a pelagic tunicate (Pyrosoma atlanticum). Isme Commun 1, 11. doi: 10.1038/s43705-021-00007-1.

Tilstone, G., Figueiras, F., Fermín, E., and Arbones, B. (1999). Significance of nanophytoplankton photosynthesis and primary production in a coastal upwelling system (Ría de Vigo, NW Spain). Mar Ecol Prog Ser 183, 13–27. doi: 10.3354/meps183013.

Turner, J. T. (2015). Zooplankton fecal pellets, marine snow, phytodetritus and the ocean’s biological pump. Prog Oceanogr 130, 205–248. doi: 10.1016/j.pocean.2014.08.005.

Vargas, C. A., Martínez, R. A., Cuevas, L. A., Pavez, M. A., Cartes, C., González, H. E., et al. (2007). The relative importance of microbial and classical food webs in a highly productive coastal upwelling area. Limnol Oceanogr 52, 1495–1510. doi: 10.4319/lo.2007.52.4.1495.

Verity, P., and Smetacek, V. (1996). Organism life cycles, predation, and the structure of marine pelagic ecosystems. Mar Ecol Prog Ser 130, 277–293. doi: 10.3354/meps130277.

Vilgrain, L., Maps, F., Picheral, M., Babin, M., Aubry, C., Irisson, J., et al. (2021). TraitLJbased approach using in situ copepod images reveals contrasting ecological patterns across an Arctic ice melt zone. Limnol Oceanogr 66, 1155–1167. doi: 10.1002/lno.11672.

Ware, D. M., and Thomson, R. E. (2005). Bottom-Up Ecosystem Trophic Dynamics Determine Fish Production in the Northeast Pacific. Science 308, 1280–1284. doi: 10.1126/science.1109049.

Weber, E. D., Auth, T. D., Baumann-Pickering, S., Baumgartner, T. R., Bjorkstedt, E. P., Bograd, S. J., et al. (2021). State of the California Current 2019–2020: Back to the Future With Marine Heatwaves? Frontiers Mar Sci 8, 709454. doi: 10.3389/fmars.2021.709454.

Wilkerson, F. P., Dugdale, R. C., and Barber, R. T. (1987). Effects of El Niño on new, regenerated, and total production in eastern boundary upwelling systems. J Geophys Res Oceans 92, 14347– 14353. doi: 10.1029/jc092ic13p14347.

Williams, P. J. le B., and Ducklow, H. W. (2019). The microbial loop concept: A history, 1930–1974. J Mar Res 77, 23–81. doi: 10.1357/002224019828474359.

Williams, R. J., Howe, A., and Hofmockel, K. S. (2014). Demonstrating microbial co-occurrence pattern analyses within and between ecosystems. Front Microbiol 5, 358. doi: 10.3389/fmicb.2014.00358.

Yamazaki, H., Mackas, D. L., and Denman, K. (2002). “Coupling small-scale physical processes with biology,” in Biological-physical interactions in the sea., eds. A. R. Robinson, J. J. McCarthy, and and B. J. Rothschild (John Wiley and Sons), 51–112.

Zamora-Terol, S., Mckinnon, A. D., and Saiz, E. (2014). Feeding and egg production of Oithona spp. in tropical waters of North Queensland, Australia. J Plankton Res 36, 1047–1059. doi: 10.1093/plankt/fbu039.

